# The chaperone Jjj2 regulates nucleoporin condensation in budding yeast

**DOI:** 10.64898/2026.07.22.740058

**Authors:** A Agote-Aran, P Gallardo, R Mancini, J S Fischer, T Bergsma, F Montesi, L Lucius, K Keuenhof, E Lorentzon, F Uliana, J L Höög, LM Veenhoff, K Weis

## Abstract

Nuclear pore complexes (NPCs) mediate nucleocytoplasmic transport through a selective permeability barrier established by nucleoporins (Nups) that contain intrinsically disordered phenylalanine-glycine (FG) repeats. Due to these domains, FG-Nups are prone to condensation, with a potential to transition into insoluble aggregates. How cells keep FG-Nups in a soluble, functional state during NPC assembly and within the native NPC remains poorly understood. Here, we identify the uncharacterized yeast J-domain protein Jjj2 as a Nup chaperone. Disrupting Jjj2 function or its interaction with Hsp70 chaperones triggers the accumulation of newly synthesized Nups in cytoplasmic condensates. Conversely, Jjj2 overexpression suppresses Nup condensation but also disrupts the NPC permeability barrier and is highly toxic. Overall, our data show that Jjj2, in concert with Hsp70, controls Nup phase state. This activity must be tightly regulated; while Jjj2 prevents condensation of newly produced Nups, its overactivity compromises nucleocytoplasmic compartmentalization.

## INTRODUCTION

The Nuclear Pore Complex (NPC) is a 60-120 MDa protein assembly that mediates the transport of biomolecules between the nucleus and the cytoplasm in all eukaryotic cells (Petrovic et al., 2025; Gall, 1954). This channel-shaped complex is embedded within the nuclear envelope (NE) and facilitates the passive diffusion of small molecules while regulating the active transport of larger cargoes. This active process is regulated by nuclear transport receptors (NTRs) that recognize specific nuclear localization or export signals (NLS and NES, respectively) in their cargo molecules (Mobbs et al., 2026). The NPC is composed of approximately 30 distinct proteins called nucleoporins (Nups) which are present in multiple copies. These Nups organize into subcomplexes that assemble with eight-fold rotational symmetry to form the scaffold inner and outer rings and asymmetric appendages, including the cytoplasmic export platform and the nuclear basket (Fig. 1A) (Lin and Hoelz, 2019; Beck and Hurt, 2017). Approximately one-third of these Nups contain phenylalanine-glycine (FG) repeats (Denning et al., 2003) and are thus termed FG-Nups. These FG repeats are natively unfolded, intrinsically disordered regions (IDRs) that create the selective permeability barrier of the NPC (Fig. 1A) (Frey et al., 2006; Wente and Rout, 2010).

**FIGURE 1:**
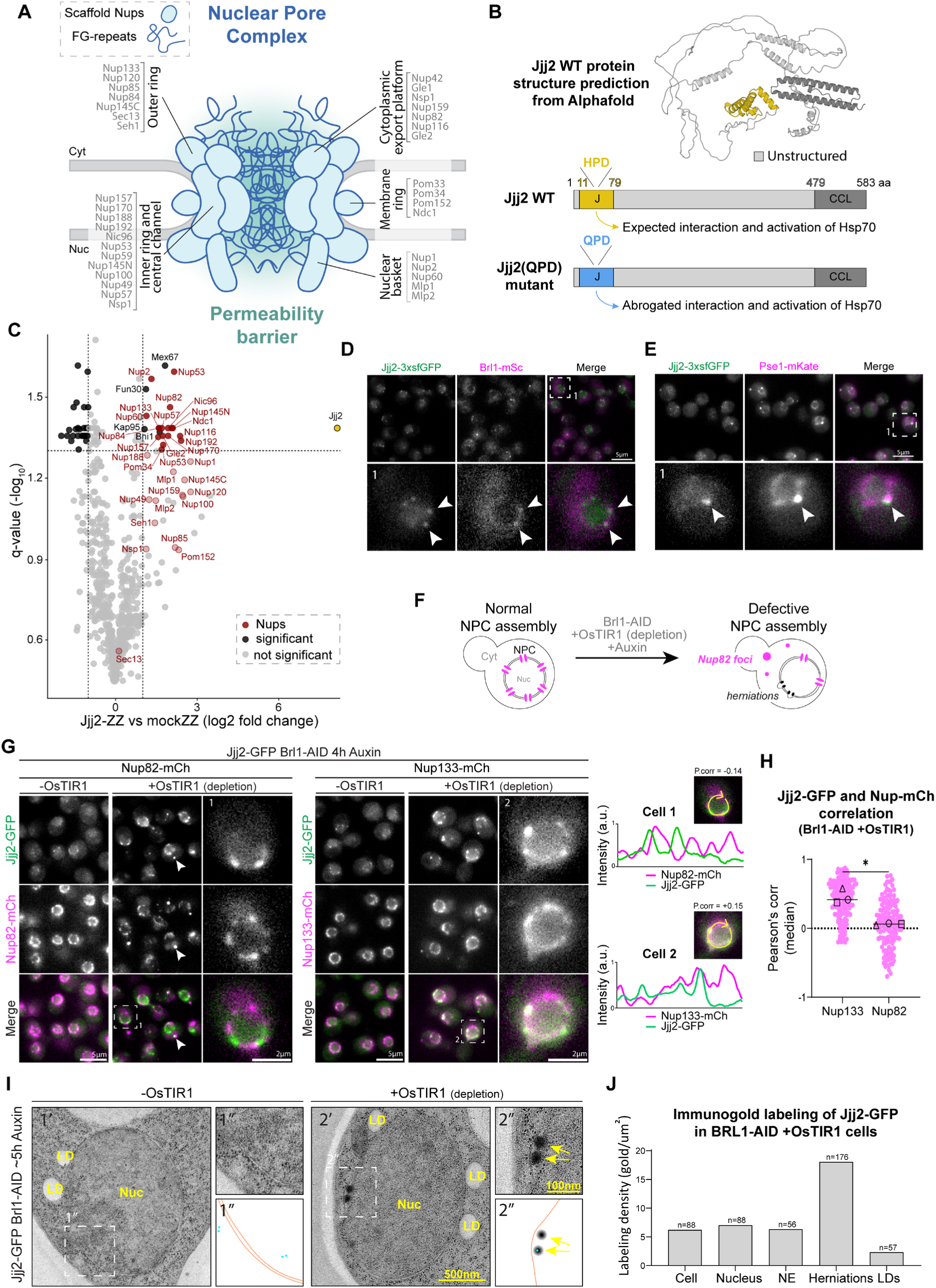
Jjj2 interacts with Nups and localizes to NPC assembly intermediates. **(A)** Schematic of the NPC architecture. Subcomplexes are labeled in black, and individual Nups in grey. Blue round shapes represent scaffold Nups; blue lines represent intrinsically disordered FG-regions. **(B)** Jjj2 structural organization. Top: AlphaFold2 structure prediction of Jjj2. The J-domain (J) is shown in yellow, the coiled-coil-like (CCL) domain in dark grey and predicted intrinsically disordered regions (low pLDDT score) in light grey. Bottom: domain architecture of Jjj2 WT indicating amino acid positions and color-coded as in the structure prediction. The HPD-to-QPD mutation in the J-domain is known to abolish its ability to interact with and stimulate the ATPase activity of Hsp70 chaperones while preserving substrate binding capacity. **(C)** Volcano plot showing the differential protein abundance in affinity purifications of ZZ-tagged Jjj2 versus soluble ZZ-tag alone. Points represent the mean log2 fold change (n=3 replicates). Significance (q-value) was calculated with the two-tailed Student’s t-test with Benjamini–Hochberg multiple comparison correction. Nups are highlighted in red and Jjj2 is highlighted in yellow. Dark colors indicate significant hits, and light colors non-significant hits. Nup hits are indicated in dark or light red color. **(D-E)** Representative fluorescence micrographs of WT cells expressing Jjj2-3xsfGFP and either Brl1-mScarlet3 or Pse1-mKate (bona fide NPC assembly factors). White arrowheads indicate foci at the NE where Jjj2 colocalizes with Brl1 or Pse1, representing physiological assembly intermediates. Scale bar: 5 μm. **(F)** Schematic of the Brl1-AID depletion system. Acute depletion of Brl1-AID is induced by auxin addition in cells expressing the E3 ligase OsTIR1 (+OsTIR1). Brl1 depletion halts NPC biogenesis, leading to the accumulation of NE herniations. Nup82-mCherry localizes to all NPCs in control conditions but is excluded from assembly intermediates upon Brl1 depletion, remaining only at mature NPCs and cytoplasmic condensates. **(G)** Left: Representative fluorescence micrographs of Brl1-AID depleted cells expressing Jjj2-GFP and either Nup82-mCherry or Nup133-mCherry. White arrowheads indicate colocalization in cytoplasmic condensates. Zoom panels (1 and 2) show Jjj2-GFP enrichment at the NE; Jjj2 signal anti-correlates with Nup82-mCherry (mature NPCs) but overlaps with Nup133-mCherry (mature NPCs and nascent intermediates). Right: Line profiles of fluorescence intensity along the NE (marked by yellow arrows) for the indicated cells. P. corr indicates Pearson’s correlation coefficient. **(H)** Pearson’s correlation coefficient analysis of the fluorescence signals at the NE from cells in (E). Pink dots represent individual cells; black shapes indicate the median of each biological replicate (n=3). 50 cells were quantified per condition, per replicate. Statistical analysis: unpaired t-test with Welch’s correction; * = pval<0.05 **(I)** Immunogold electron micrographs showing Jjj2-GFP (gold particles) in Brl1-AID cells approximately 5 h after auxin addition. Brl1-depleted cells show electron-dense NE herniations (yellow arrows). Bottom right: schematic drawings of the micrographs; orange lines represent the NE and blue dots represent Jjj2-GFP associated gold particles. **(J)** Gold particle labeling density (particles/μm^2^) across different cellular compartments. Lipid droplets (LDs) serve as a negative control. The number of quantified compartments is indicated above each bar.

Due to their disordered nature and ability to weakly self-interact, FG-Nups have the tendency to undergo phase separation. Purified FG-Nups can phase-separate into liquid or gel-like condensates that recapitulate some of the properties of the permeability barrier (Celetti et al., 2020; Frey et al., 2006; Frey and Görlich, 2007; Schmidt and Görlich, 2015). However, over time or in conditions with increased crowding, FG-Nups can also form amyloid fibrils (Ader et al., 2010; Milles et al., 2013). In a cellular context, various conditions, such as defects in NPC biogenesis (Kralt et al., 2022; Lone et al., 2015; Makio et al., 2009; Scarcelli et al., 2007; Zhang et al., 2018b), the overexpression of FG-Nup levels (Boer et al., 1998; Thomas et al., 2023), mislocalization of Nup translation (Lautier et al., 2021) or different stress conditions (Thomas et al., 2023; Zhang et al., 2018a), trigger the formation of cytoplasmic Nup condensates that recruit NTRs and other Nups. Notably, Nup overexpression and the presence of these cytoplasmic foci correlate with compromised growth and toxicity in human cell cultures and *C. elegans* (Boer et al., 1998; Thomas et al., 2023) suggesting that tight regulation of Nup condensation is crucial for cell survival.

Recent studies have begun to elucidate how cells regulate Nup homeostasis during their journey from initial synthesis at the ribosome to final incorporation into the NPC (Kuiper et al., 2023; Penzo and Palancade, 2023; Veldsink et al., 2023). During this transit, Nups are particularly vulnerable to aberrant interactions or inappropriate phase separation in the cytoplasm. Consequently, cells have evolved diverse mechanisms to maintain Nups in an assembly-competent state. For instance, the localized translation of Nups at the NPC (Lautier et al., 2021) and the co-translational assembly of specific Nup subcomplexes serve to minimize inappropriate Nup-Nup interactions that could otherwise interfere with proper NPC assembly (Lautier et al., 2021; Seidel et al., 2021). Furthermore, NTRs act as specialized chaperones by binding Nups, sometimes co-translationally, to keep them soluble until they are released at the nuclear side during NPC assembly (Lautier et al., 2021; Ryan et al., 2007; Seidel et al., 2021; Walther et al., 2003). Accordingly, Importin β has been shown to prevent FG-Nup aggregation *in vitro* (Milles et al., 2013; Otto et al., 2024). Beyond NTRs, classical molecular chaperones have also been shown to play a critical role in actively regulating Nup phase state in mammalian cells. *In vitro* and in human cells, DNAJB6b (a J-domain protein) or a ternary complex consisting of DNAJB6b, Hsp70 and MLF2 maintains FG-Nup condensates in a functional state, preventing their progression into irreversible aggregates (Bergsma et al., 2026, 2025; Kuiper et al., 2022; Prophet et al., 2022).

In the present work, we sought to uncover novel Nup regulators implicated in the early stages of NPC biogenesis in budding yeast. This approach built upon our characterization of the temporal order of NPC assembly (Onischenko et al., 2020; Kralt et al., 2022) to identify proteins that bind transiently to early assembly intermediates but dissociate upon structural maturation. By screening for factors that preferentially enrich with early-assembling Nups rather than mature NPCs, several putative Nup regulators were identified, with the J-domain protein Jjj2 emerging as a primary candidate for further investigation.

Jjj2 is a protein of unknown function that, like DNAJB6b, belongs to the diverse family of J-domain proteins of which there are 21 members in *S. cerevisiae* (Ruger-Herreros et al., 2024; Kampinga and Craig, 2010). J-domain proteins function as essential cofactors for Hsp70 chaperones, mediating substrate recognition and driving the activation of Hsp70 ATPase activity to facilitate vital cellular tasks such as protein disaggregation, decondensation, and structural remodeling (Rosenzweig et al., 2019). The evolutionarily invariant HPD motif within the J-domain is strictly required for the interaction with and activation of Hsp70 (Kityk et al., 2018; Tsai and Douglas, 1996).

In this study, we show that Jjj2 is a chaperone that interacts with newly produced Nups to regulate their condensation state. In the absence of Jjj2, or upon mutation of its HPD motif, a subset of Nups forms cytoplasmic condensates, suggesting that both Jjj2 and an Hsp70 activity are required for this regulatory function in cells. We show that Jjj2 modulates Nup condensates reducing their size and intensity and preventing their transition into aggregates. Interestingly, increased Jjj2 levels induce NPC permeability defects and are toxic for yeast cells, indicating that Jjj2 can also influence the phase state of FG-Nups within the native NPC.

## RESULTS

### Proteomic screening identifies the J-domain protein Jjj2 as a putative Nup regulator

In our previous work, we characterized the interphasic NPC assembly pathway in budding yeast using the KARMA method to categorize nucleoporins (Nups) into early, intermediate, and late assembly tiers (Onischenko et al., 2020). Building on these findings, we sought to identify additional non-Nup factors involved in the early stages of NPC biogenesis. We hypothesized that such regulators associate transiently with assembling Nups along their journey from translation to pore integration but are not part of the mature NPC. To identify such factors, we took advantage of the KARMA proteomic dataset to find proteins enriched in pulldowns of early-tier Nups relative to the late-tier marker Mlp1. This approach already successfully identified the transmembrane protein Brl1 as an NPC assembly factor (Kralt et al., 2022), which had been previously identified and verified also by others (Lone et al., 2015; Saitoh et al., 2005; Zhang et al., 2021, 2018b).

The present study expanded this investigation by characterizing four additional high-ranking candidates identified by this screening strategy (Fig. S1A; Suppl. Table 1). Along with Brl1 (included as a positive control), we selected: Jjj2, a J-domain protein of unknown function; Gpi17 and Gpi16, components of the GPI transamidase complex residing at the ER (Harris and Marles-Wright, 2024); and Nmd3, a ribosomal subunit export factor (Trotta et al., 2003).

To examine a potential role of these candidates in NPC assembly or Nup regulation, we utilized an auxin-inducible degron (AID) system for their acute depletion (Fig. S1B). As a readout for Nup condensation and NPC assembly defects, we monitored the localization of Nup82, which is known to accumulate in cytoplasmic condensates when it cannot be incorporated at the NPC during impaired assembly (Kralt et al., 2022; Lone et al., 2015; Makio et al., 2009; Scarcelli et al., 2007; Zhang et al., 2018b). While Brl1 depletion induced cytoplasmic Nup82 foci in ∼70% of cells, Jjj2 depletion showed the strongest effect among the new candidates (∼30% of cells), leading us to focus on Jjj2 (Fig. S1C-E).

### Jjj2 interacts with Nups and localizes to NPC assembly intermediates

Jjj2 is a low-abundance J-domain protein in budding yeast with an estimated copy number of 542 +/- 427 molecules per cell ((Ho et al., 2018) dataset accessed through the Saccharomyces Genome Database). AlphaFold2 structural predictions (Jumper et al., 2021; Varadi et al., 2024, 2022) indicate that Jjj2 contains two folded domains comprising approximately 30% of the sequence: an N-terminal J-domain and a C-terminal coiled-coil-like region. The remaining 70% of the protein exhibits long stretches of low confidence scores (pLDDT<70), suggesting this region is likely intrinsically disordered (Fig. 1B, Fig. S1F).

To confirm the interaction between Jjj2 and Nups observed in the KARMA assay (Fig. S1A; Suppl. Table 1), we performed an affinity purification coupled to mass spectrometry on endogenously tagged Jjj2. Nups belonging to all NPC subcomplexes were enriched in Jjj2 pulldowns, supporting an interaction between Jjj2 and Nups (Fig. 1C; Suppl. Table 2).

Next, we investigated the subcellular localization of Jjj2 in wild-type (WT) cells by tagging the endogenous protein with a triple superfolder GFP (Jjj2-3xsfGFP). Jjj2-3xsfGFP displayed a diffuse distribution throughout the cell with a slight enrichment in the nucleoplasm, while also localizing to a few distinct foci at the NE (Fig. 1D-E). Because Jjj2 was restricted to these sparse puncta rather than continuously decorating the entire NE rim (a pattern characteristic of mature NPCs), we reasoned that these foci might represent NPC assembly intermediates. Supporting this hypothesis, Jjj2 colocalized at these foci with *bona fide* NPC assembly factors, such as Brl1 and Pse1 (Kralt et al., 2022; Lusk et al., 2002; Zhang et al., 2018b) (Fig. 1D-E).

To determine whether Jjj2 is also recruited to stalled NPC assembly sites, we analyzed its localization in mutants where NPC biogenesis is arrested. To this end, we used the acute Brl1-AID depletion system (Fig. 1F), which triggers the formation of stalled assembly intermediates visible as NE herniations by electron microscopy (Kralt et al., 2022). These herniations incorporate Nup133 but not Nup82; consequently, the NE-associated Nup82 signal is restricted entirely to pre-existing NPCs that matured before Brl1 depletion while unincorporated Nup82 accumulates in cytoplasmic condensates (Fig. 1F).

Following Brl1 depletion, Jjj2 accumulated in prominent NE foci that anti-correlated with the Nup82 signal characteristic of mature NPCs. Instead, these Jjj2 foci overlapped with Nup133, which marks both assembly intermediates and mature pores (Fig. 1G). Pearson’s correlation analysis confirmed that Jjj2 associates preferentially with defective assembly intermediates (Nup133-positive/Nup82-negative) and is excluded from mature NPCs (Nup133-positive/Nup82-positive) (Fig. 1H). Interestingly, Jjj2 also localizes to the cytoplasmic Nup82 condensates (Fig. 1G, arrowheads), suggesting that it also interacts with the cytoplasmic pool of Nups in addition to NPC assembly intermediates.

To confirm that Jjj2 localizes to stalled NPC assembly intermediates, we performed immunogold electron microscopy. In Brl1-depleted cells, Jjj2-associated gold particles were significantly enriched within electron-dense NE herniations compared to other cellular structures (Fig. 1I-J). Consistent with these findings, Jjj2 also localized to defective NPCs in several other assembly mutants (*gle2Δ, nup116Δ*, pMet-NUP170 *nup157Δ*, *nup120Δ*, *apq12Δ, nup188Δ, nup133Δ*) (Aitchison et al., 1995; Heath et al., 1995; Li et al., 1995; Makio et al., 2009; Murphy et al., 1996; Scarcelli et al., 2007; Wente and Blobel, 1993) (Fig. S2A-D).

Collectively, these ultrastructural and fluorescence data demonstrate that Jjj2 selectively localizes to physiological assembly intermediates, defective NPCs, and cytoplasmic Nup condensates, while being excluded from mature, fully functional NPCs.

### Jjj2 deletion triggers the formation of cytoplasmic Nup condensates

Given the broad interaction between JJJ2 and Nups in our affinity purification, we investigated whether the deletion of Jjj2 affects the localization of Nups belonging to different subcomplexes of the NPC (Fig. 1A). Central channel FG-Nups were excluded from this analysis, as their fluorescent tagging compromises cellular fitness (Otto et al., 2024). While total protein levels and NE intensities of Nups remained unchanged (Fig. S3A-B, Fig. 2A-B), several Nups (Nup192, Nup116, Nup159, and Nup82) formed cytoplasmic foci (Fig. 2A, C). Nup116 and Nup159 contain FG-repeats and they form a heterotrimer with Nup82 on the cytoplasmic side of the NPC (Yoshida et al., 2011). Nup192 is a scaffold Nup and structurally similar to the karyopherin family (Stuwe et al., 2014). These Nup foci were sensitive to 1,6-hexanediol (1,6-HD) treatment (Fig. S3C-D), an aliphatic alcohol that disrupts weak hydrophobic interactions which are usually involved in condensate formation. Because more solid gel-like condensates are generally less susceptible to 1,6-HD, this sensitivity is consistent with the foci being in a relatively liquid-like state.

**FIGURE 2:**
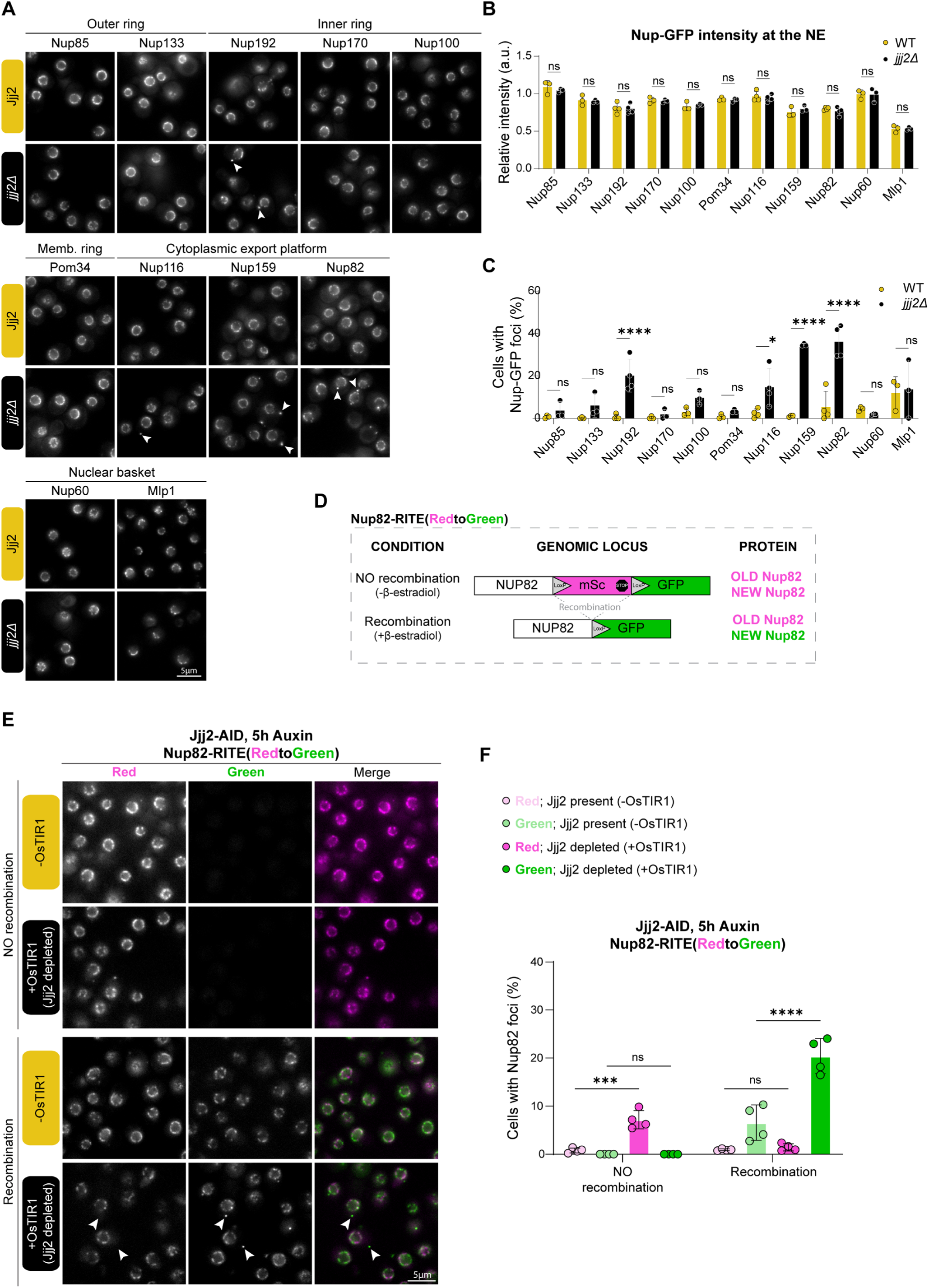
Jjj2 is required for the correct localization of newly produced Nups. **(A)** Representative fluorescence micrographs of GFP-tagged Nups in wild-type (Jjj2) and Jjj2-deletion (*jjj2Δ*) cells. Scale bar: 5 μm. **(B)** Quantification of Nup-GFP fluorescence intensity at the nuclear envelope (NE) from cells in (D). Intensities are normalized to the average intensity of the Y-complex (Nup85 and Nup133) in the WT background. Data represent mean ±SD from at least three biological replicates. Statistical test: Two-way ANOVA with Sidak’s multiple comparison test. *p < 0.05, **p < 0.01, ***p < 0.001, ****p < 0.0001. **(C)** Quantification of the percentage of cells in (D) showing cytoplasmic Nup-GFP foci. Data represent the mean ±SD from at least three biological replicates. Statistical test: Two-way ANOVA with Sidak’s multiple comparison test. *p < 0.05, **p < 0.01, ***p < 0.001, ****p < 0.0001. **(D)** Schematic of the Recombination-Induced Tag Exchange (RITE) strategy. This system enables the visualization of pre-existing (“old,” mScarlet3; red) and newly produced (“new,” GFP; green) Nup82 populations. Recombination is mediated by β-estradiol-induced Cre recombinase acting on *LoxP* sites. The genomic configuration and the corresponding protein products are indicated for both non-induced (-β-estradiol) and induced (+β-estradiol) conditions. **(E)** Recombination was induced with (+β-estradiol) for 5.5 h to switch expression from “old” Nup82-mScarlet3 to “new” green Nup82-GFP; non-induced cells (-β-estradiol) served as NO recombination control. Jjj2-AID was acutely depleted with auxin 30 min post-recombination (+OsTIR1) for a duration of 5 h. Control cells (-OsTIR1) express Jjj2 at endogenous levels. At this time point, both Nup82 populations form a distinct nuclear envelope rim, enabling clear discrimination of cytoplasmic foci from mature NPCs. Arrowheads indicate cytoplasmic foci positive for newly produced Nup82-GFP and negative for pre-existing Nup82-mScarlet3. Scale bar: 5 μm. **(F)** Percentage of cells showing Nup82 cytoplasmic foci in the mScarlet3 (“old”; pink bars) and GFP (“new”; green bars) channels. Data correspond to the 5.5 h time point described in (B). Mean ±SD from at least three biological replicates. Statistics: Two-way ANOVA; *** = pval<0.001, **** = pval<0.0001.

Notably, *jjj2Δ* cells exhibited no growth defects (Fig. S3E), even when challenged with heat shock or a prolonged stationary phase (Fig. S3F-G). Furthermore, electron microscopy revealed no structural NPC abnormalities or NE herniations characteristic of NPC assembly mutants such as Brl1-depleted cells (Fig. S3H).

Together, these observations suggest that Jjj2 is required to prevent the condensation of specific Nups, but cell growth or NPC assembly are not obviously impaired in the absence of Jjj2.

### Newly produced Nups form the cytoplasmic condensates in *jjj2Δ* cells

What is the origin of the cytoplasmic Nup condensates observed in *jjj2Δ* cells? Because Jjj2 localizes to the Nup82 condensates that accumulate during NPC assembly defects (Fig. 1G), and given that these specific foci are composed of newly made Nups (Kralt et al., 2022), we hypothesized that the condensates in *jjj2Δ* mutants similarly originate from the cytoplasmic pool of nascent, unincorporated Nups. To test this hypothesis, we used the Recombination-Induced Tag Exchange (RITE) method, which enables the differential visualization of recently translated (“new”) versus pre-existing (“old”) protein populations (Verzijlbergen et al., 2010).

We tagged endogenous Nup82 with a tandem fluorescent RITE cassette (Nup82-RITE(RedtoGreen)) consisting of a LoxP-flanked mScarlet3 followed by a downstream GFP. Under steady-state conditions, Nup82-mScarlet3 is constitutively expressed; upon induction of β-estradiol-inducible Cre recombinase, the mScarlet3 gene is excised, triggering a permanent switch to Nup82-GFP expression (Fig. 2D). This transition allows for spatial and temporal discrimination between the “old” red Nup82 and the “new” green Nup82 populations in the cell.

Quantification revealed that Jjj2 depletion triggered a significant increase in the percentage of cells bearing “new” Nup82-GFP condensates, whereas the “old” Nup82-mScarlet3 population remained unaffected (Fig. 2E-F). Notably, the two fluorescent tags displayed different baseline propensities to form condensates, with Nup82-GFP exhibiting a higher baseline frequency of foci compared to Nup82-mScarlet3. Regardless of this baseline variation, the relative increase in condensation following Jjj2 depletion occurred exclusively within the newly synthesized Nup82-GFP pool (Fig. 2F). Therefore, these results show that Jjj2 regulates the condensation of newly made Nups.

### Jjj2 requires Hsp70 chaperone activity in yeast cells

We next sought to determine Jjj2’s role within the broader chaperone network and whether its function requires Hsp70 partners.

In the canonical Hsp70 cycle, a J-domain protein recognizes specific substrates and delivers them to its Hsp70 partner, subsequently stimulating Hsp70’s ATPase activity to drive downstream chaperone functions. To test whether Jjj2 operates through this classical mechanism, we mutated its conserved J-domain HPD motif to QPD. This HPD to QPD mutation is widely established to preserve substrate binding while abolishing the JDP’s ability to interact with and activate Hsp70 (Fig. 1B) (Tsai and Douglas, 1996; Zhang et al., 2023). We replaced the endogenous *JJJ2* gene with the *jjj2(QPD)-3xHA* mutant allele and assessed the subcellular localization of Nup82-GFP. Quantification of cells displaying cytoplasmic Nup82-GFP foci revealed that the Jjj2(QPD) mutation phenocopies the *jjj2Δ* deletion (Fig. 3A-B). Expectedly, given that *jjj2Δ* cells grow normally, the Jjj2(QPD) mutation had no impact on baseline cell growth (Fig. S4A). These results demonstrate that Jjj2’s role in preventing Nup condensation depends on its functional coupling with an Hsp70 partner. Furthermore, localization analysis of this mutant tagged with 3xsfGFP revealed that while Jjj2(QPD) still associated with occasional NE foci like the wild-type protein, it was also strongly colocalized with the Nup82 cytoplasmic foci (Fig. 3C, arrowheads), suggesting this mutant can still interact with the newly produced Nup82 but cannot regulate its condensation state.

**FIGURE 3:**
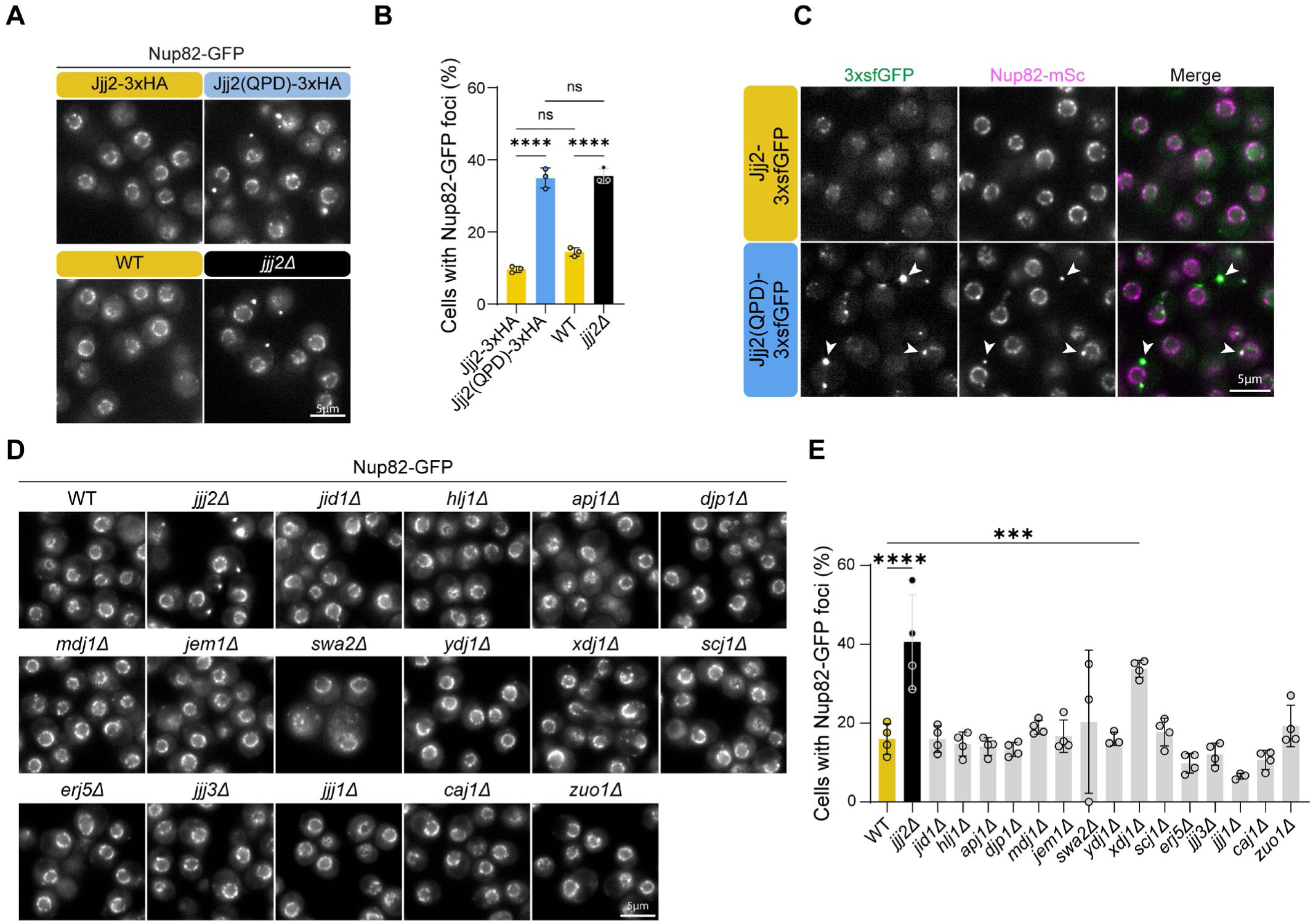
Jjj2 requires Hsp70 chaperone activity in cells and displays functional selectivity toward Nups among yeast J-domain proteins. **(A)** Representative fluorescence micrographs of Nup82-mScarlet3 co-expressed with either Jjj2-3xsfGFP or Jjj2(QPD)-3xsfGFP. White arrowheads indicate the colocalization of Jjj2(QPD) and Nup82-mScarlet3 in cytoplasmic condensates. **(B)** Representative fluorescence micrographs of Nup82-GFP in the indicated genetic backgrounds: Jjj2-3xHA, Jjj2(QPD)-3xHA, untagged WT Jjj2, and *jjj2Δ*. Scale bar: 5 μm. **(C)** Percentage of cells from (C) exhibiting Nup82-GFP cytoplasmic foci. Data represent mean ±SD from at least three biological replicates. Statistical test: One-way ANOVA with Sidak’s multiple comparison test. ****p < 0.0001. **(D)** Representative fluorescence micrographs of Nup82-GFP in cells lacking the indicated non-essential J-domain protein genes. Scale bar: 5 μm. **(E)** Percentage of cells from (E) exhibiting Nup82-GFP cytoplasmic foci. Data represent mean ±SD from at least three biological replicates. Statistical test: One-way ANOVA with Sidak’s multiple comparison test. ***p < 0.001, ****p < 0.0001. Note: *swa2Δ* cells exhibit slower growth, consistent with previously published literature (Du et al., 2001; Martineau et al., 2010). Due to the limited number of cells available for quantification in this specific strain, the data exhibit a wider standard deviation.

### Jjj2 displays functional selectivity toward Nups among yeast J-domain proteins

Given the extensive redundancy of the yeast chaperone network (Gong et al., 2009), we asked if other J-domain proteins act alongside Jjj2 to regulate Nup condensation. We systematically deleted all non-essential J-domain proteins in a Nup82-GFP background (Fig. 3D). While the majority of deletions failed to trigger cytoplasmic Nup82 foci, loss of Xdj1 (*xdj1Δ*) induced foci that were qualitatively smaller than those observed in *jjj2Δ* cells (Fig. 3D-E).

However, a *jjj2Δ xdj1Δ* double mutant (as well as other J-domain protein double-deletion combinations) showed no synergistic increase in Nup82 condensation or growth defects (Fig. S4B-D). These findings show that, among J-domain proteins, Jjj2 exhibits distinct functional selectivity towards Nups among yeast J-domain proteins. Furthermore, the absence of a fitness phenotype in *jjj2Δ* or its corresponding double mutants may suggest that the redundant network safeguarding Nup homeostasis may be complex, likely involving multiple J-domain proteins or overlapping roles with distinct chaperone classes.

### Jjj2 modulates Nup condensation in cells

Our findings suggest that Jjj2 promotes the soluble state of newly synthesized Nups. To directly test if Jjj2 modulates the phase state of Nups, we utilized a recently established system (Bergsma et al., 2025; Otto et al., 2024) utilizing a galactose-inducible, FG-rich region of Nup100 (pGAL1-GFP-Nup100FG). This construct lacks the structured domain required for NPC scaffold interaction and accumulates in cytoplasmic and nucleoplasmic particles. Because GFP-Nup100FG overexpression is proteotoxic, we induced expression for 2 hours with galactose (2h time point) followed by the addition of glucose and a 4-hour recovery period (6h time point) (Fig. 4A). Particle frequency and their sizes and shapes were then quantified using PhaseMetrics (Bergsma et al., 2025).

**FIGURE 4:**
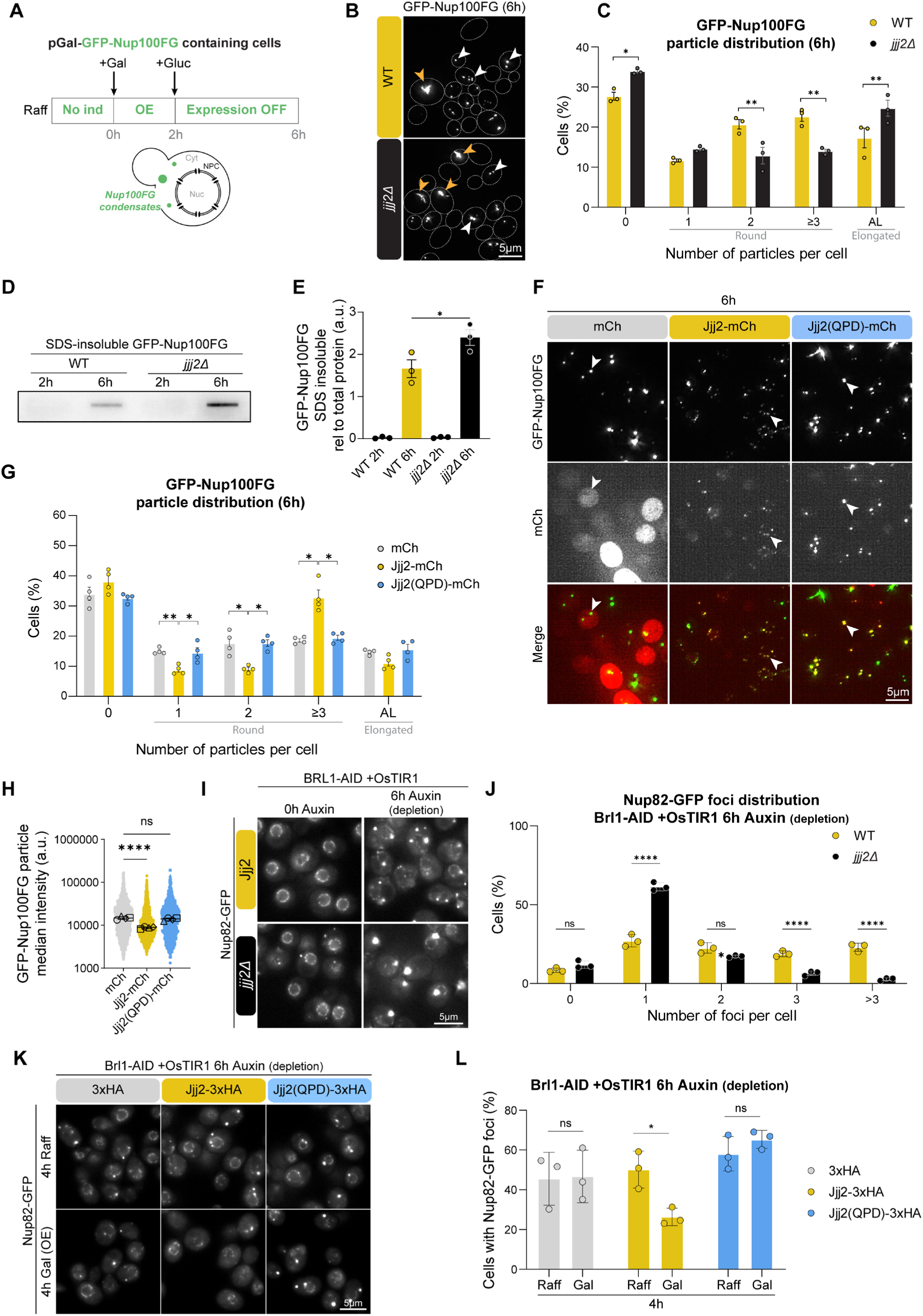
Jjj2 modulates Nup condensates. **(A)** Schematic of the GFP-Nup100FG condensate inducing system. The FG-rich region of Nup100 (Nup100FG), N-terminally tagged with GFP, is inducibly expressed via galactose addition for 2 h (“OE”; 2 h time point). Expression is subsequently repressed by the addition of glucose for a 4 h recovery period (“Expression OFF”; 6 h time point). **(B)** Representative fluorescent micrographs of WT and *jjj2Δ* cells at the 6h time point according to (A). White arrowheads indicate round liquid/gel-like condensates; orange arrowheads indicate elongated aggregate-like structures of GFP-Nup100FG. Scale bar: 5 μm. **(C)** Percentage of cells from (B) categorized by the number of round condensates (0, 1, 2, or ≥3) or the presence of elongated aggregate-like (AL) structures. Data represent mean ±SD from three biological replicates. Statistical test: two-way ANOVA with Sidak’s multiple comparison test. *p < 0.05, **p < 0.01. **(D)** Filter trap assay of WT and *jjj2Δ* cells at the 6h time point according to (A) to detect the 0.5% SDS-resistant fraction of GFP-Nup100FG normalized to the total GFP-Nup100FG protein levels (Fig. S5A-B). **(E)** Band intensities on filter trap assays in (D) are normalized to total protein levels. Data represent mean ±SD from three biological replicates. Statistical test: one-way ANOVA with Dunnett’s multiple comparison test. *p < 0.05. **(F)** Representative fluorescence micrographs of cells co-overexpressing either mCherry (mCh) control, Jjj2-mCh, or Jjj2(QPD)-mCh together with GFP-Nup100FG at the 6 h time point according to (A). White arrowheads indicate round condensates. **(G)** Quantification of the distribution of cells in (F) categorized by the number of round condensates (0, 1, 2, or ≥3) or the presence of elongated AL structures. Data represent the mean ±SD from four biological replicates. Statistical test: two-way ANOVA with Sidak’s multiple comparison test. *p < 0.05, **p < 0.01. **(H)** Superplot showing the median fluorescence intensity of GFP-Nup100FG particles from (F). Individual dots represent single particles; larger black circles indicate the median of each biological replicate (n=3). Statistical test: one-way ANOVA with Dunnett’s multiple comparison test. ****p < 0.0001. **(I)** Representative fluorescence micrographs of Nup82-GFP in WT or *jjj2Δ* cells with Brl1-AID depletion for 0 h or 6 h. Scale bar: 5 μm. **(J)** Percentage of cells from (I) categorized by the number of Nup82-GFP condensates per cell (0, 1, 2, 3, or >3). Data represent mean ±SD from three biological replicates. Statistical test: two-way ANOVA with Sidak’s multiple comparison test. ****p < 0.0001. **(K)** Representative fluorescence micrographs of Nup82-GFP in cells upon Brl1-AID depletion (6 h) combined with overexpression (Gal) of 3xHA (control), Jjj2-3xHA, or Jjj2(QPD)-3xHA for the final 4 h. Raffinose (Raff) serves as the non-overexpression control. Scale bar: 5 μm. **(L)** Percentage of cells from (K) showing Nup82-GFP condensates. Data represent mean ±SD from three biological replicates. Statistical test: two-way ANOVA with Sidak’s multiple comparison test. *p < 0.05.

The Nup100FG construct typically forms a heterogeneous population of particles: round liquid/gel-like condensates and elongated, aggregate-like structures (Fig. 4B; white and orange arrowheads, respectively) (Bergsma et al., 2025). Deletion of JJJ2 resulted in a significant increase in the percentage of cells with aggregate-like particles and particle-free cells, with a concomitant decrease in cells with two or more round condensates (Fig. 4B-C). This suggests an accelerated maturation of GFP-Nup100FG into stable aggregates in the absence of Jjj2, with particle-free cells likely resulting from the asymmetric inheritance of large aggregates by the mother cell. Filter trap assays (FTA) confirmed that the SDS-insoluble fraction of GFP-Nup100FG significantly increased in *jjj2Δ* cells at the 6h time point (Fig. 4D-E, Fig. S5A-B). Such aggregates were less frequent at the 2h time point in the microscopy experiments (Fig. S5C) or not detected in the FTA (Fig. 4D-E), further supporting the idea that they result from condensate maturation over time. This GFP-Nup100FG SDS insoluble fraction likely represents amyloid aggregates, since the Nup100 FG-region has been previously reported to form amyloid-like aggregates (Danilov et al., 2023; Halfmann et al., 2012).

Given these results, we wanted to test whether increasing Jjj2 levels would decrease Nup100FG condensation. To test this, we co-overexpressed either Jjj2 WT or Jjj2(QPD) with Nup100FG. Overexpression of WT Jjj2 reduced both the size and fluorescence intensity of Nup100FG condensates and increased the number of condensates per cell (shifting the population toward ≥3 condensates) (Fig. 4F-H), suggesting a failure of smaller foci to coalesce and/or big foci being dispersed into smaller foci. Notably, Jjj2 colocalized with GFP-Nup100FG condensates and aggregates, suggesting that it can interact with the FG-repeat regions. In contrast, while Jjj2(QPD) still colocalized with the Nup100FG foci, it failed to reduce their size or intensity (Fig. 4F-H) confirming that the activity of Jjj2 depends on the interaction with Hsp70. Moreover, Jjj2 appears to exhibit a certain degree of selectivity toward Nup substrates rather than acting as a generic anti-condensation factor. When co-overexpressed with the human protein TDP-43, an unrelated, condensation-prone protein, Jjj2 failed to colocalize with TDP-43 foci or alter their distribution, frequency and intensity (Fig. S5D-F).

We then assessed the effect of Jjj2 on more physiological Nup condensates. For this, we used the Brl1-AID depletion system, which is characterized by an accumulation of cytoplasmic Nup82 condensates (Fig. 1F). While the deletion of JJJ2 did not affect Nup82 levels (Fig. S5G) or further increase the overall percentage of cells containing Nup82 condensates compared to WT (Fig. S5H), it significantly altered the distribution of condensate frequency per cell (Fig. 4I-J). In cells with WT background, we observed a relatively even distribution of cells containing one to four condensates. In contrast, *jjj2Δ* cells exhibited a larger proportion of cells containing only one focus (Fig. 4J). These foci appeared larger than those in WT cells (Fig. 4I), suggesting that Nup82 condensates undergo increased coalescence or reduced dispersion in the absence of Jjj2. In addition, we examined whether Jjj2 overexpression could rescue the condensation phenotype. Notably, overexpression of WT Jjj2, but not the Jjj2(QPD) mutant, significantly decreased the percentage of cells exhibiting Nup82 condensates (Fig. 4K-L; Fig. S5I-J).

### Jjj2 directly modulates Nup condensation *in vitro*

To determine if Jjj2 directly interacts with FG-Nups, we purified recombinant Nup100FG and Jjj2 proteins and incubated them in a buffer containing 10% PEG, a condition previously established to induce Nup100FG condensation (Bergsma et al., 2025). In line with the *in vivo* findings, Jjj2 full length (FL) significantly reduced the fraction of SDS-insoluble Nup100FG aggregates (Fig. S6A-B) and decreased condensate size and intensity (Fig. S6C-F). This effect required Jjj2’s C-terminal region, as Jjj2(aa1-132), a truncated version lacking most of the predicted intrinsically disordered region and the C-terminal coiled-coil-like region, did not show these capabilities. These results indicate that Jjj2 can directly interact with Nup100FG and, at high concentrations, it is biochemically sufficient to modulate Nup100FG condensate properties and reduce Nup100FG aggregation in an Hsp70-independent manner.

### Jjj2 overexpression leads to NPC permeability barrier defects and reduces cell viability

In our Jjj2 overexpression experiments, we observed that while Nup condensation was decreased, the cells exhibited strong growth defects. Spotting assays confirmed that overexpression of WT Jjj2 is highly toxic, whereas the Jjj2(QPD) mutant showed only a minimal growth defect compared to the control strain (Fig. 5A). This suggests that an excess of Jjj2’s Hsp70-dependent activity severely compromises cell fitness.

**Figure 5:**
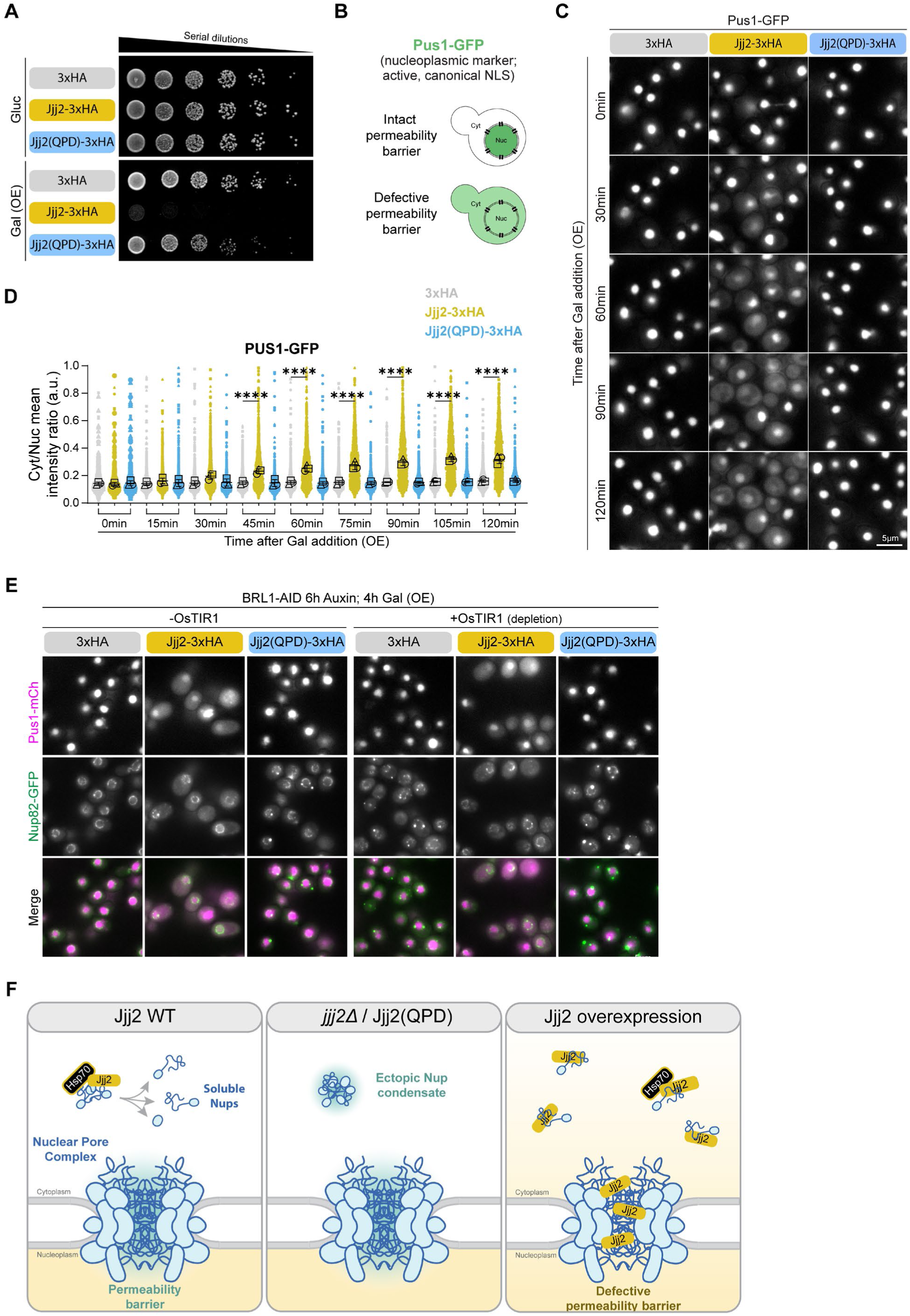
Jjj2 overexpression leads to permeability barrier defects and decreases cell viability. **(A)** Spotting assay of wild-type cells expressing 3xHA, Jjj2-3xHA or Jjj2(QPD)-3xHA from the GAL1 promoter. Cells were grown on medium containing either glucose (expression OFF) or galactose (expression ON). **(B)** Schematic of the nucleoplasmic marker Pus1-GFP. In an intact permeability barrier, Pus1-GFP is exclusively nuclear; a defective barrier allows passive leakage into the cytoplasm. **(C)** Representative fluorescence micrographs from a time-lapse experiment monitoring Pus1-GFP localization in the indicated time points upon overexpression of 3xHA, Jjj2-3xHA, or Jjj2(QPD)-3xHA. Scale bar: 5 μm. **(D)** Superplot showing the Pus1-GFP cytoplasmic/nucleoplasmic mean intensity ratio from cells in (C) including additional time points. Small colored shapes represent individual cells; larger black shapes indicate the mean of each biological replicate (n=3). Statistical test: one-way ANOVA with Tukey’s multiple comparison test. ****p < 0.0001. **(E)** Representative fluorescent micrographs showing Pus1-mCh and Nup82-GFP localization in cells with (+OsTIR1) or without (-OsTIR1) Brl1-AID depletion for 6 h with auxin. 3xHA, Jjj2-3xHA or Jjj2(QPD)-3xHA expression was induced with galactose addition 2 h after auxin addition (galactose present during the last 4 h of the experiment). Note that Pus1-mCherry leaks even when new NPC assembly is halted. Scale bar: 5 μm. **(F)** Proposed model for Jjj2 function. Jjj2 acts as a dedicated chaperone for FG-Nups. Under physiological conditions, Jjj2 interacts with newly produced cytoplasmic FG-Nups to compete with inter-FG hydrophobic interactions, thereby decreasing the FG-Nup tendency to form condensates or decondensing already formed FG-Nup condensates. Hsp70 activity is required for Jjj2 release, enabling “chaperone cycling” to manage the pool of newly synthesized Nups. In *jjj2Δ* or Jjj2(QPD) cells, unchaperoned FG-Nups form cytoplasmic condensates. Conversely, Jjj2 overexpression leads to non-specific interactions with the FG-meshwork within the central channel of mature NPCs, weakening the permeability barrier and resulting in cellular toxicity.

We reasoned that increased levels of Jjj2 might act on the endogenous FG-Nup meshwork within the central channel and compromise the nucleocytoplasmic compartmentalization which is essential for viability. To test this, we monitored the localization of Pus1, an endogenous nucleoplasmic protein containing multiple classical Lys/Arg-rich nuclear localization signal (NLS) motifs previously used as a nuclear marker (Romanauska and Köhler, 2018; Wang et al., 2024) (Fig. 5B). In time-lapse imaging experiments, Pus1 was initially restricted to the nucleus but progressively leaked into the cytoplasm starting approximately 45 minutes after the induction of Jjj2 WT overexpression (Fig. 5C-D; Fig. S7A-D). This mislocalization was not observed in the Jjj2(QPD) mutant, indicating that the leakage and toxicity phenotypes correlate and are Hsp70-dependent.

To determine whether this compartmentalization defect stemmed from disrupted active transport or a permeability barrier failure, we evaluated two additional transport reporters (Fig. S7A). First, we tested an active transport reporter consisting of a blue fluorescent protein (BFP) fused to the strong NLS of RPL25 (RPL25NLS-BFP; (Schaap et al., 1991)). Unlike with Pus1-GFP, the nuclear accumulation of RPL25NLS-BFP was barely altered upon Jjj2 overexpression (Fig. S7E-G), indicating that the active import pathway of this reporter remained functional, though reporter-specific differences in nuclear retention cannot be formally ruled out. Second, we monitored the passive reporter MGM4 (a ∼230 kDa GFP-maltose binding protein fusion), which lacks an NLS and is normally excluded from the nucleus due to its large size (Popken et al., 2015). Consistent with a permeability barrier defect, MGM4 mislocalized to the nucleus upon Jjj2 WT overexpression but remained cytoplasmic in the Jjj2(QPD) mutant (Fig. S7H-J). In line with a selective barrier defect rather than a fundamental collapse of the active transport, Jjj2 overexpression did not globally disrupt NTRs’ localization as many retained their steady-state distribution (Fig. S8A).

We next investigated whether these permeability barrier defects resulted from structural damage to the NPC or the NE. Quantitative fluorescence microscopy revealed no changes in Nup intensities at the NE, suggesting that NPC stoichiometry and number remain stable within the timeframe of Jjj2 overexpression (Fig. S9A-B). Ultrastructural analysis by electron microscopy did not reveal NE ruptures, indicating that the compartmentalization defect is not resulting from discontinuities in the NE (Fig. S9C). Finally, we addressed whether the observed permeability barrier defects arise from errors in NPC assembly or whether already existing NPCs are affected. To this end, we depleted Brl1 to arrest *de novo* NPC biogenesis prior to inducing Jjj2 expression. Whereas no leakage could be detected upon Brl1 depletion alone, Jjj2 overexpression still triggered nuclear leakage even when NPC assembly was blocked (Fig. 5E). This indicates that the Jjj2-induced permeability defects are caused by the disruption of mature NPCs.

Collectively, these results show that an excess of Jjj2 compromises the permeability barrier of fully assembled NPCs. Jjj2 likely alters the biophysical integrity of the FG-meshwork in the NPC channel, resulting in a more permissive barrier that increases passive diffusion across the NE. The observed mislocalization of transport reporters is likely a kinetic phenomenon: active import can compensate for a leaky barrier if the import rate exceeds the rate of passive diffusion. However, for proteins with slower import kinetics or passive cargoes, the barrier defect leads to a loss of compartmentalization.

## DISCUSSION

In this study, we identified Jjj2 as a putative NPC assembly factor or Nup regulator under the assumption that such factors preferentially interact with early-assembling Nups compared to late-assembling ones (Fig. S1A; Suppl. Table 1). Under wild-type conditions, Jjj2 localizes to NPC assembly intermediates (Fig. 1D-E), and it is strongly enriched at stalled intermediates that form in assembly-deficient mutants (Fig. 1G-J; Fig. S2A-D). Functionally, the acute depletion or genetic deletion of Jjj2 leads to the mislocalization of newly synthesized Nups in cytoplasmic foci (Fig. 2A, C).

Mechanistically, Jjj2 modulates the phase state of Nups. Cells show increased Nup condensates and aggregates upon Jjj2 deletion (Fig. 4A-E, I-J), whereas Jjj2 overexpression decreases condensate number and size (Fig. 4F-H, K-L). Notably, this anti-condensation function requires Hsp70 chaperone activity *in vivo*, as the J-domain mutant Jjj2(QPD) phenocopies the *jjj2Δ* null mutant (Fig. 3A-C). However, Hsp70 remains dispensable for Jjj2-mediated decrease in Nup condensation and aggregation *in vitro* (Fig. S6A-F).

Intriguingly, an excess of WT Jjj2 is highly toxic to yeast cells (Fig. 5A), a phenotype that correlates with a defect in the permeability barrier and an increase in passive diffusion across the NE (Fig. 5C-D; Fig. S7B-J). Moreover, Jjj2 overexpression triggers nuclear leakage even when *de novo* NPC biogenesis is blocked by Brl1 depletion (Fig. 5E). These findings indicate that the permeability barrier defects do not arise from misassembled NPCs, but rather from the disruption of the permeability barrier in mature NPCs.

Based on these data, we propose a model (Fig. 5F) where Jjj2 interacts with newly produced Nups to prevent their aberrant condensation or aggregation, potentially by physically competing with inter-molecular FG-FG interactions from FG-Nups, which are known to be important to establish the permeability barrier (Frey et al., 2006; Beck and Hurt, 2017). Within the cell, the Hsp70 ATPase cycle drives Jjj2-Nup dissociation; this chaperone recycling allows a relatively small pool of Jjj2 molecules to sequentially regulate a more abundant population of condensation-prone Nups. Given the lack of enrichment of ribosomal components in our Jjj2 affinity purifications (Fig. 1C), we do not favor a model that Jjj2 engages with its substrates co-translationally, but instead hypothesize that Jjj2 binds to Nups shortly after their synthesis and accompanies them during their transit to assembling pore sites, abandoning the complex once the NPC reaches structural maturity. In this manner, we envision Jjj2 as a “true chaperone” that accompanies Nups from birth to integration into the mature NPC, ensuring that they avoid non-productive interactions and phase separation along the way.

Our proposed model raises several questions about how Jjj2 recognizes its clients and functions within the cellular environment. A central question is how Jjj2 selectively targets newly produced Nups while avoiding mature NPCs. Although a definitive mechanism remains beyond the scope of this manuscript, it would be intriguing to investigate the basis of this selectivity in future studies. For instance, this selectivity could rely on localization-dependent post-translational modifications on either Jjj2 or its Nup clients. Furthermore, it remains to be fully determined whether Jjj2 primarily acts upstream to maintain baseline Nup solubility or actively dismantles pre-formed condensates. Finally, the fate of the aberrant Nup condensates that accumulate in the absence of Jjj2, whether they are eventually cleared by autophagy, targeted by protein quality control pathways, or solubilized by alternative means, is an open question that our current data cannot answer.

Intriguingly, despite the apparent importance of maintaining Nup solubility for proper pore assembly, *jjj2Δ* cells displayed no baseline growth or survival defects (Fig. S3E-G). This lack of a phenotypic consequence could stem from functional redundancy with other quality control mechanisms and/or the broader chaperone network (Gong et al., 2009), potentially involving distinct chaperone classes beyond the J-domain proteins tested here. Alternatively, Jjj2 may act as a conditional factor required only under specific stresses that challenge Nup homeostasis. While acute heat shock and quiescence did not alter *jjj2Δ* cell survival (Fig. S3F-G), Jjj2’s functional importance might manifest under alternative physiological challenges yet to be explored.

A paradox in our findings is that Hsp70 activity is required for Jjj2 function *in vivo* (Fig. 3A-C) but dispensable *in vitro* (Fig. S6A-F). This discrepancy could be explained by an intracellular stoichiometric imbalance: Jjj2 is a low-abundance protein in yeast cells, whereas our *in vitro* condensation assays were conducted at an equimolar ratio (Fig. S6A-F). We hypothesize that Jjj2 interacts with Nup100FG via its unstructured C-terminal region through weak hydrophobic interactions that compete with inter-FG contacts. *In vitro*, abundant Jjj2 can saturate FG-regions through direct binding to reduce phase separation. However, in the cellular environment, where Jjj2 is limiting and aggregation-prone Nups vastly outnumber it, Hsp70-mediated ATP hydrolysis is likely required to release Jjj2 from these engagements and promote chaperone recycling. This recycling amplifies the operational capacity of Jjj2, allowing a small pool of Jjj2 molecules to sequentially protect a much larger substrate population. Notably, however, elevating Jjj2 expression *in vivo* does not bypass this Hsp70 dependency. Whether this overexpressed pool remains limiting relative to the total cellular Nup burden, or if other complex intracellular factors strictly enforce this *in vivo* requirement, remains to be determined.

Jjj2 appears to be a functional yeast counterpart of human DNAJB6b. Although structurally distinct, both chaperones selectively localize to NPC assembly intermediates but not mature NPCs, and directly regulate FG-Nup phase separation (Bergsma et al., 2026; Kuiper et al., 2022; Prophet et al., 2022). Mechanistically, DNAJB6b contains a serine/threonine (S/T)-rich domain, which also contains phenylalanine residues, that is critical for binding FG-repeats and amyloid-like substrates (Bergsma et al., 2026; Kakkar et al., 2016; Kuiper et al., 2022). By contrast, Jjj2 lacks such an obvious S/T-rich region. Our data show that Jjj2’s anti-condensation capacity resides within its unstructured C-terminal region (aa 132–583), as the truncated Jjj2(aa1-132) variant cannot modulate Nup100FG condensation (Fig. S6A-F).

Importantly, while DNAJB6b interacts with a broad spectrum of amyloidogenic clients (including Huntingtin, FUS, TDP-43, and α-synuclein) (Kakkar et al., 2016; Resnick et al., 2025; Udan-Johns et al., 2014), Jjj2 seems to exhibit a certain client selectivity toward Nups. This selectivity is highlighted by its lack of colocalization with, or effect on, heterologously co-expressed TDP-43 condensates (Fig. S5D-F). Nonetheless, a broader panel of alternative substrates must be evaluated to map the full endogenous client repertoire of Jjj2, including whether it can modulate functional or pathogenic amyloid-like structures native to yeast, such as prions.

Finally, the nuclear permeability barrier defects triggered by Jjj2 overexpression provide intriguing insight into the biophysical nature of the NPC transport barrier. Our data argue against the notion that an excess of Jjj2 leads to a disruption of NPC assembly (Fig. 5E), altered pore stoichiometry (Fig. S9A-B), NE ruptures (Fig. S9C), or a fundamental collapse of active transport (Figs. 5C-D, S7B-G). Rather, our findings suggest that an excess of Jjj2 compromises the permeability barrier by directly infiltrating and altering the organization of the FG-Nups within the central channel of the NPC, thereby disrupting its intrinsic barrier integrity and lowering the threshold for passive passage. As such, Jjj2 emerges as a tool capable of modulating the internal environment of the NPC *in vivo*.

In summary, leveraging the preferential interaction of Nup regulators with early-assembling over late-assembling Nups has proven to be an effective approach for identifying novel players in NPC homeostasis, as demonstrated here by our identification of the Nup condensation regulator Jjj2. Future studies will be essential to fully understand the precise mechanism by which Jjj2 regulates the condensation state of newly produced Nups, as well as to uncover the specific physiological conditions under which this function becomes critical for cell function and survival.

## ACKNOWLEDGMENTS

The authors would like to thank members of the Weis lab for their discussions and comments on the manuscript, especially Sarah Khawaja, Elisa Dultz, Matthias Wojtynek, Felix Räsch, Vamshidhar Gade, Katerina Radilova and Federico Cerullo. We acknowledge the Barral Lab (specifically Aliaksandr Damenikan) for kindly sharing the 3xsfGFP and Cas9 plasmids with us.

This work was supported by grants from the Swiss National Science Foundation to K.W. (TMAG-3_209354, 310030_208213, 3200-0-242936 and CRSII5_193740). A. A.-A. was supported by an EMBO Postdoctoral Fellowship (ALTF_910-2021). L. V. acknowledges the funding from the Dutch Research Council (NWO) grant no. VI.C.192.031 and OCENW.GROOT.2019.068. J.L.H. acknowledges the Swedish Science foundation (2023-04293) and Cancer foundation (21 1865 Pj) for funding.

## AUTHOR CONTRIBUTION

Conceptualization, K. W., A. A.-A and J. S. F. Investigation and formal analysis, A. A.-A., P. G., R. M., J. S. F., T. B., F. M., L. L., K. K., E. L. and F. U. Writing original draft, A. A.-A. and K.W. Writing review and editing, A. A.-A, P. G., R.M., J. S. F., T. B., E. L., F. U., J. L. H., L. V., K. W. Supervision K. W., L. V. and J. L. H. Visualization, A. A.-A., P. G., J. S. F. and T. B. Funding acquisition, K. W., A. A.-A., L. V. and J. L. H. Project administration, K. W.

## DECLARATION OF GENERATIVE AI AND AI-ASSISTED TECHNOLOGIES IN THE MANUSCRIPT

During the preparation of this manuscript, the author(s) used ChatGPT (OpenAI) and Gemini (Google) to check grammar and spelling. After, the authors reviewed and edited the content as needed and take full responsibility for the content of the published article.

## MATERIALS AND METHODS

### Data and Code Availability

The mass spectrometry proteomics data have been deposited to the ProteomeXchange Consortium via the PRIDE partner repository with the dataset identifier PXD081359.

### Plasmid and yeast strain construction

Plasmids were generated using standard molecular cloning techniques and are listed in Table 1. Budding yeast strains were constructed using standard genetic protocols; transformations and integrations of linear DNA fragments into the genome were achieved via homologous recombination.

**Table 1:**
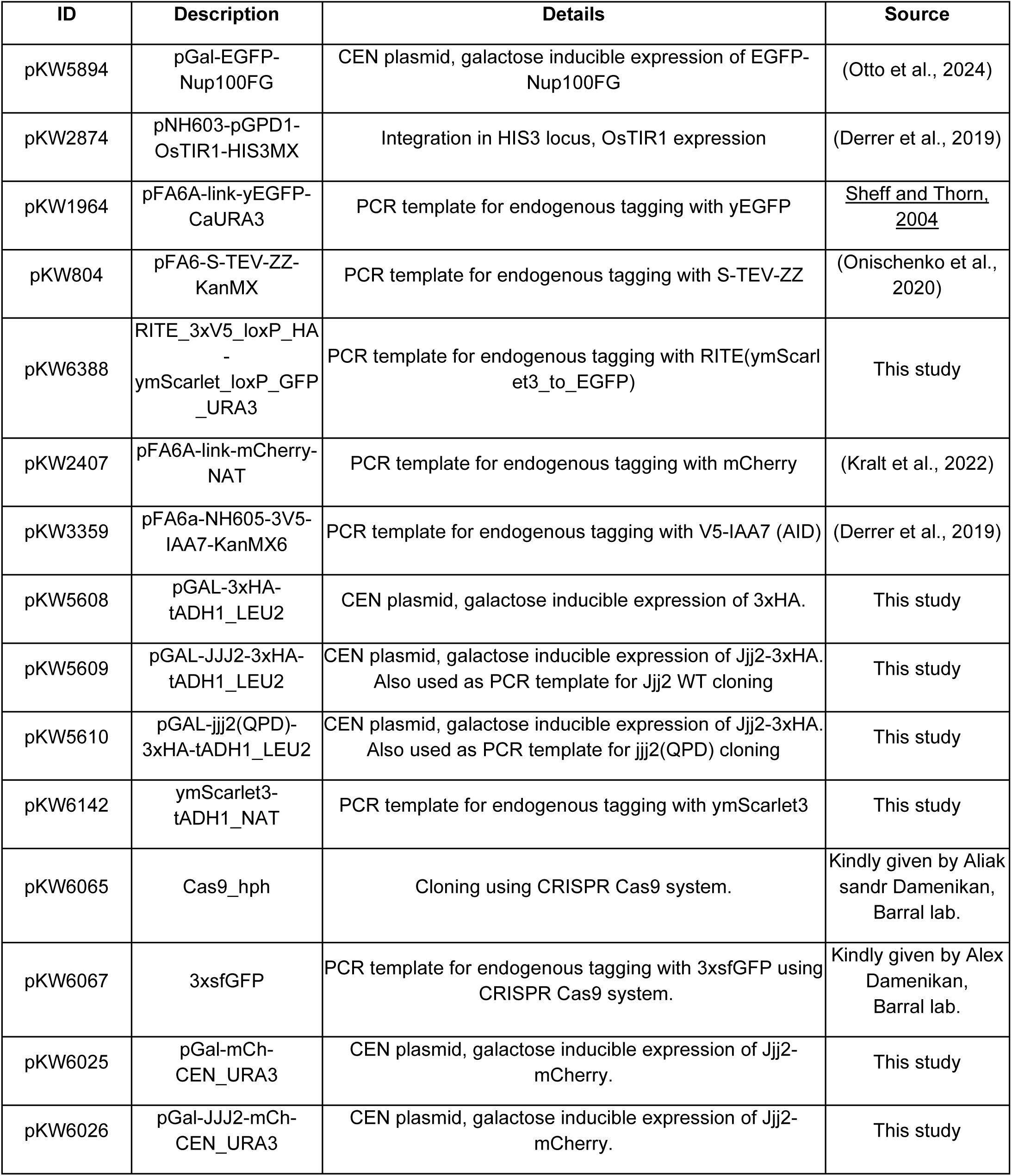
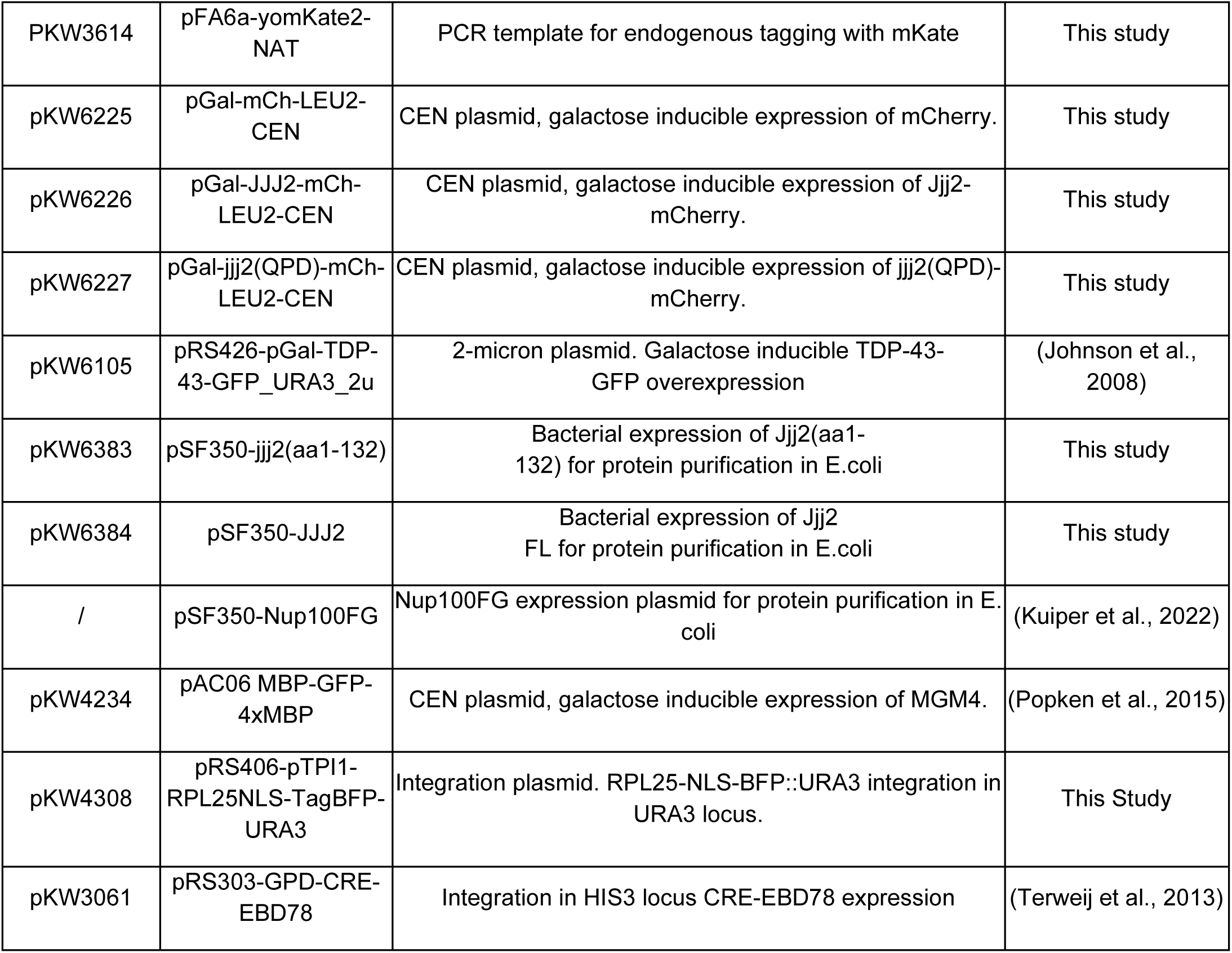
Plasmids used in this study.

Yeast strains utilized in this study are listed in Table 2. The 3xsfGFP yeast strain was generated using CRISPR-Cas9 genome editing. Detailed cloning strategies and sequence information are available upon request.

**Table 2:**
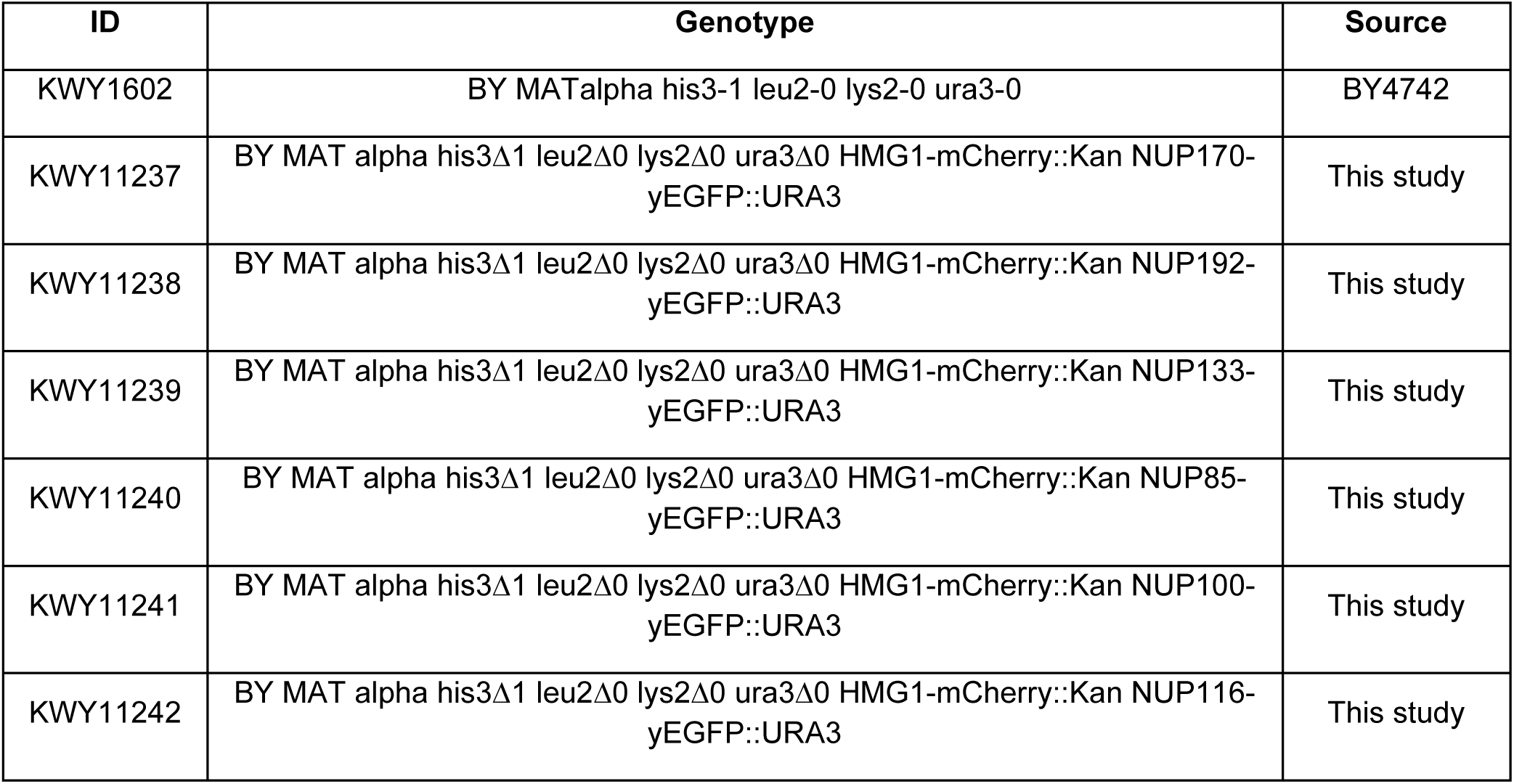
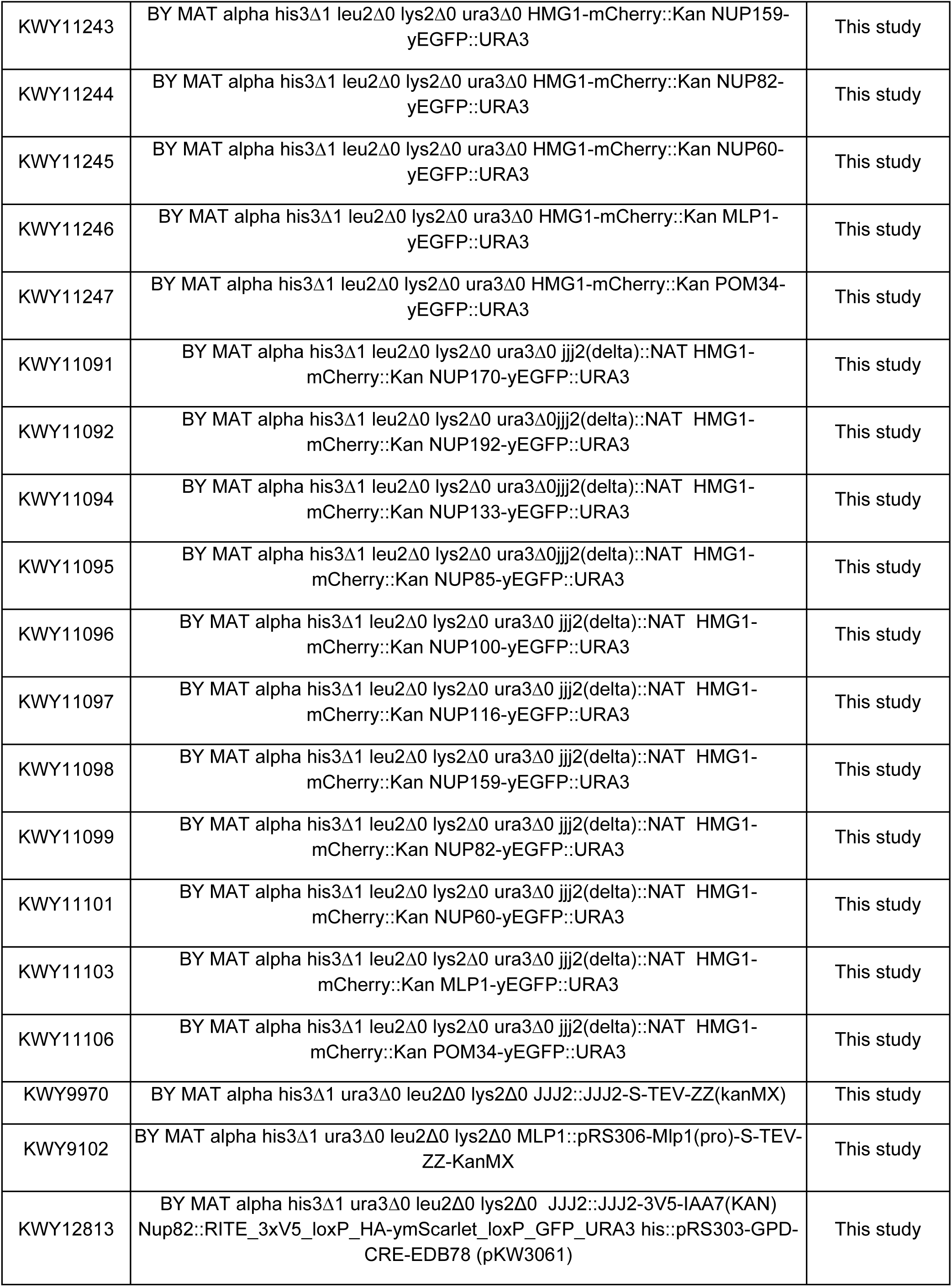
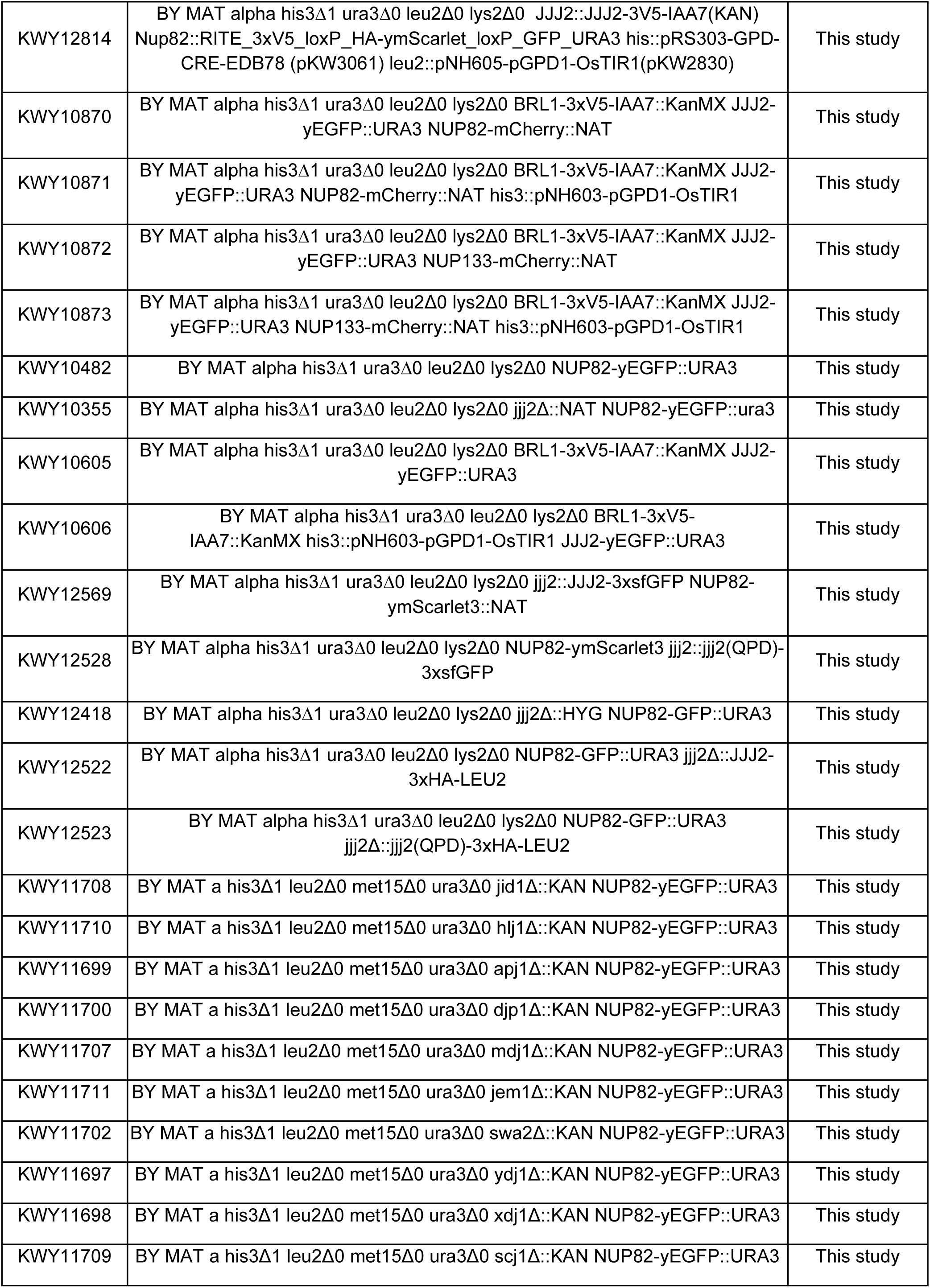
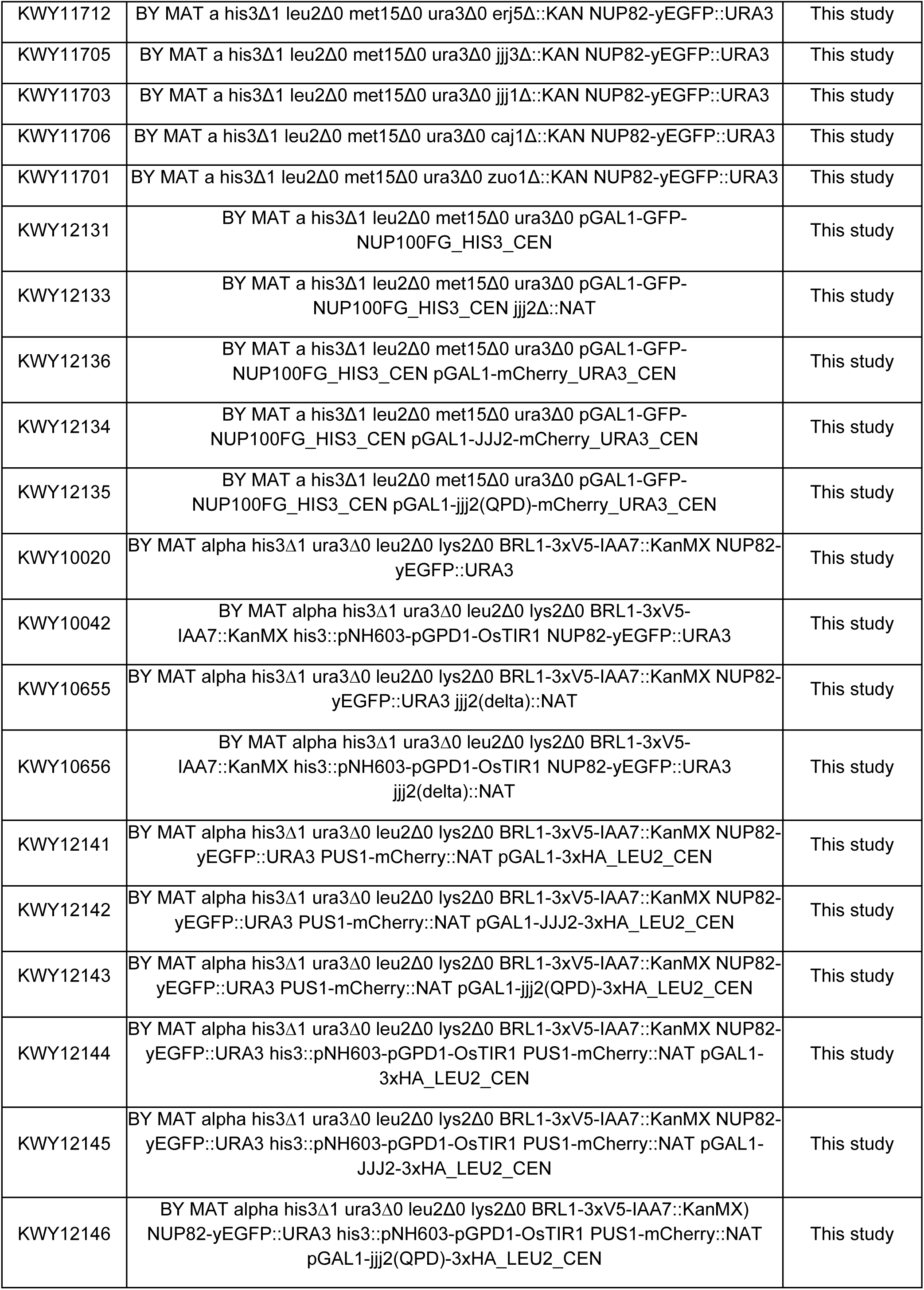
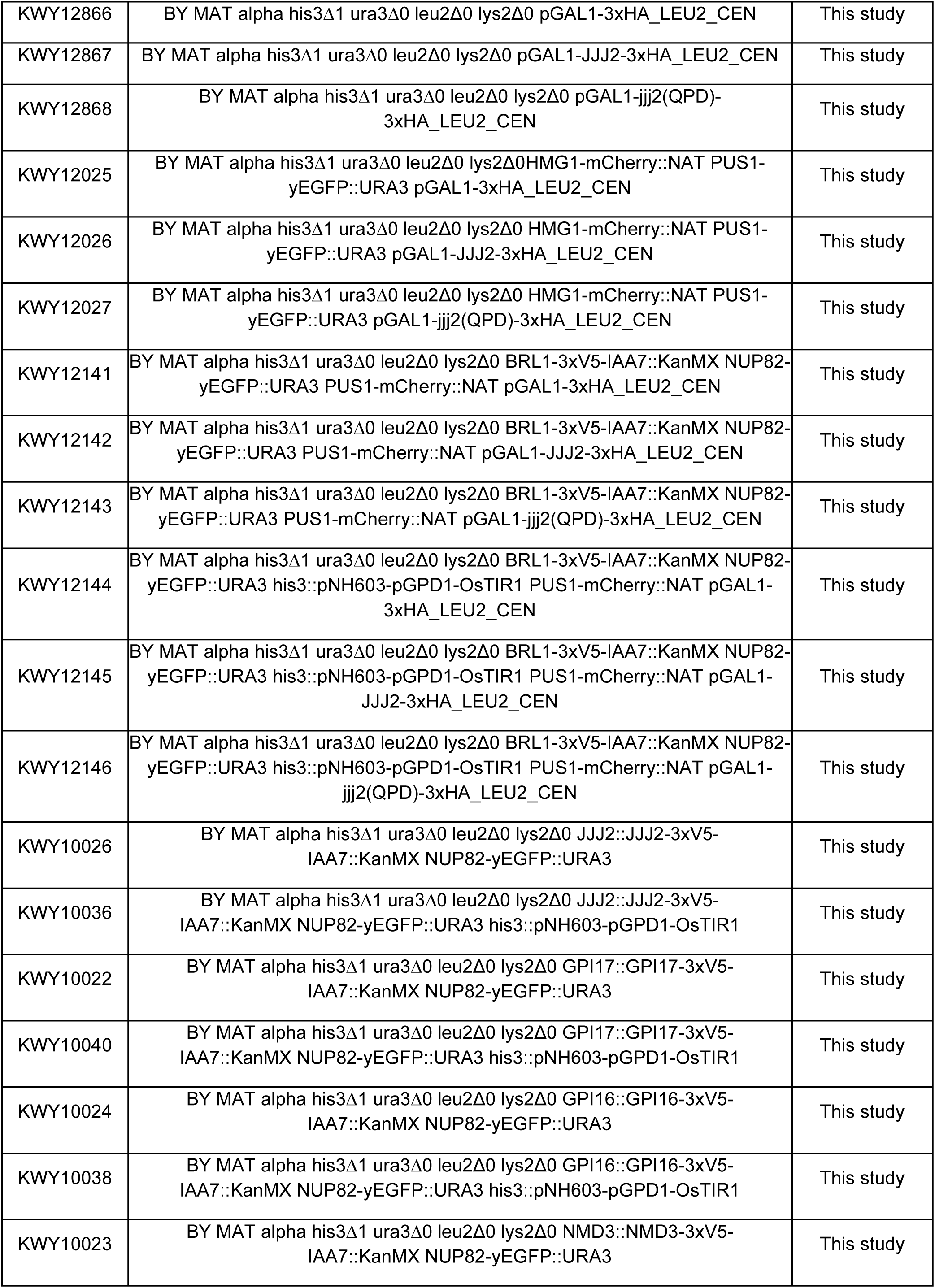
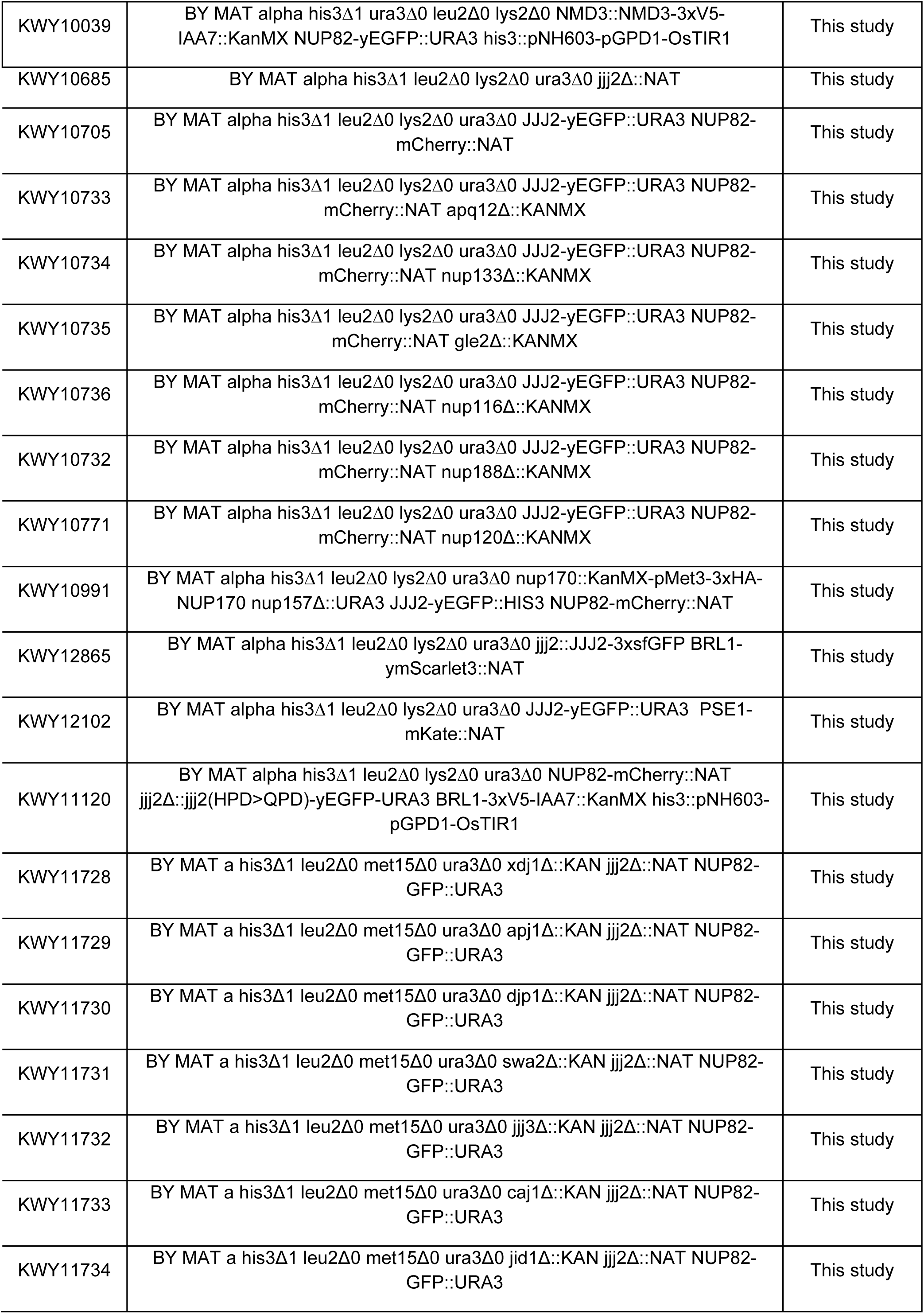
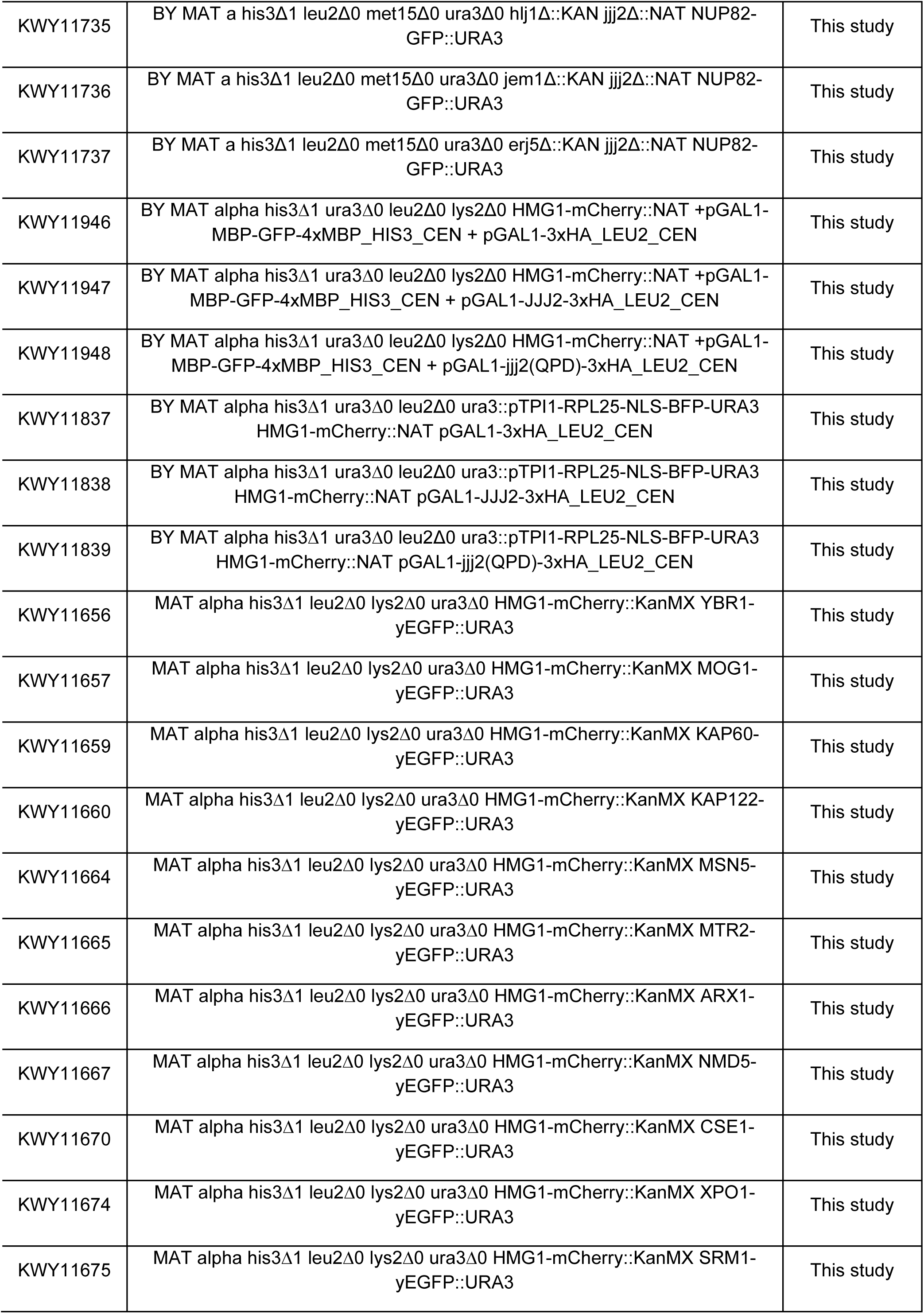
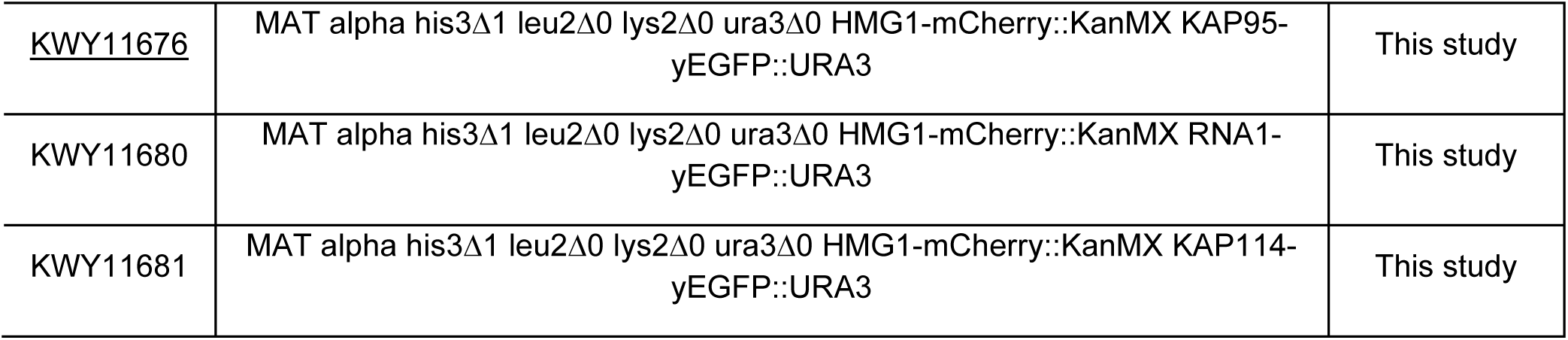
Yeast strains used in this study.

### Reagents, Antibodies and Software

All chemicals, biological reagents, antibodies, and software packages utilized in this study are indicated in Table 3.

**Table 3:** Reagents, antibodies and software.

| Reagent type | Designation | Source or reference | Identifier | Additional information / Used for |
| --- | --- | --- | --- | --- |
| Chemical compound, drug | IP6 | Sigma-Aldrich | P5681 | AID experiments |
| Chemical compound, drug | Auxin | Sigma-Aldrich | I2886 | AID experiments |
| Chemical compound, drug | β-Estradiol | Sigma-Aldrich | E8875 | RITE experiments |
| Chemical compound, drug | Galactose | Acros Organics | 150610010 | Overexpression induction |
| Chemical compound, drug | Raffinose | Chem Impex International | 2022 | Non-overexpression control |
| Chemical compound, drug | Concanavalin A (ConA) | Sigma-Aldrich | C2010 | Fluorescence microscopy |
| Chemical compound, drug | Fluorescent brightener 28 | Sigma-Aldrich | F3543 | Fluorescence microscopy |
| Other | 0.22μm MC membranes | Millipore | GSWP02500 | Immunogold/Electron microscopy |
| Chemical compound, drug | Lowicryl® HM20 Non-polar, Hydrophobic, -70°C Embedding Kit | Polysciences | 15924-1 | Immunogold/Electron microscopy |
| Chemical compound, drug | Uranyl Acetate | SPI-CHEM | 02624-AB | Immunogold/Electron microscopy |
| Chemical compound, drug | Formvar (vinylec E) | TAAB | F005 | Immunogold/Electron microscopy |
| Chemical compound, drug | Reynold's Lead citrate | Merck | Cat#109969 | Immunogold/Electron microscopy |
| Chemical compound, drug | PBS<br>(NaCl (137mM), KCl (2.7mM), Na<sub>2</sub>HPO<sub>4</sub> (10mM), KH<sub>2</sub>PO<sub>4</sub> (1.8mM) in H<sub>2</sub>O) |  |  | Immunogold/Electron microscopy |
| Chemical compound, drug | BSA | Merck (Sigma-Aldrich) | A7030 | Immunogold/Electron microscopy |
| Chemical compound, drug | Fish skin gelatin | Sigma-Aldrich | Cat# 9000-70-8 | Immunogold/Electron microscopy |
| Antibody | rb anti-GFP | Abcam | ab6556 | Immunogold/Electron microscopy |
| Antibody | goat anti rabbit, IgG 10nm gold | Electron microscopy sciences | #25108 | Immunogold/Electron microscopy |
| Chemical compound, drug | Glutaraldehyde | Sigma-Aldrich Sweden AB | Cat#3802 | Immunogold/Electron microscopy |
| Chemical compound, drug | IPTG | AppliChem | A10080025 | Protein purification |
| Chemical compound, drug | Ni Sepharose 6 FastFlow | Cytiva | 17531806 | Protein purification |
| Chemical compound, drug | Tris-HCl | Fischer Scientific | BP152-1 | Protein purification |
| Chemical compound, drug | Guanidine Hydrochloride (GuHCl) | Thermo Scientific | 24100 | Protein purification |
| Chemical compound, drug | DTT | Roche | 10708984001 | MS assays and Protein purification |
| Chemical compound, drug | Imidazole | Roth | X998.4 | Protein purification |
| Chemical compound, drug | Glycerol | Sigma-Aldrich | G5516 | Protein purification |
| Other | Vivaspin Protein Concentrator spin columns | Cytiva | 28-9322-47, 28-9322-48 | Protein purification |
| Peptide, recombinant protein | DNase I | Roche | 10104159001 | Protein purification |
| Chemical compound, drug | cOmplete Protease Inhibitor Cocktail | Roche | 5053489001 | Protein purification |
| Antibody | Primary antibody monoclonal mouse anti-GFP | Santa Cruz | sc-9996 | Filter trap assays |
| Antibody | Primary mouse monoclonal anti-tetra-His antibody | Qiagen | 34670 | Filter trap assays |
| Antibody | Secondary anti-mouse m-IgGk BP-HRP | Santa Cruz | sc-516102 | Filter trap assays |
| Other | Nitrocellulose membrane<br>Protan 0.2 µm | Amersham | GE10600001 | Filter trap assays |
| Chemical compound, drug | HEPES | Fisher Scientific | BP310-500 | Filter trap assays |
| Chemical compound, drug | Phosphate buffered saline | Sigma-Aldrich | P4417 | Filter trap assays |
| Chemical compound, drug | Tween20 | MP Biomedicals | TWEEN201 | Filter trap assays |
| Chemical compound, drug | Sodium chloride | Acros Organics | 207790010 | Filter trap assays |
| Chemical compound, drug | Magnesium chloride hexahydrate | Sigma-Aldrich | M2393 | Filter trap assays |
| Chemical compound, drug | Phenylmethanesulfonyl fluoride (PMSF) | Sigma-Aldrich | P7626 | Filter trap assays |
| Chemical compound, drug | cOmplete ULTRA tablets, Mini EDTA-free | Roche | 05892791001 | Filter trap assays |
| Chemical compound, drug | Albumine bovine serum (BSA) | Acros Organics | 268131000 | Filter trap assays |
| Chemical compound, drug | Pierce™ BCA Protein Assay Kit | Fisher Scientific | 23225 | Filter trap assays |
| Chemical compound, drug | ECL Prime Western Blotting Detection Reagent | Amersham | RPN2232 | Filter trap assays |
| Chemical compound, drug | 1.6 hexanediol | Sigma-Aldrich | 240117 | Fluorescence microscopy |
| Chemical compound, drug | Protease inhibitor cocktail | Sigma-Aldrich | P8215 | MS assays and Protein purification |
| Chemical compound, drug | Purified IgG protein from rabbit serum | Sigma-Aldrich | I5006 | MS assays |
| Chemical compound, drug | Coomassie Brilliant Blue R-250 | Bio-Rad | 161-0400 | MS assays |
| Chemical compound, drug | Sequencing grade porcine trypsin | Promega | V5113 | MS assays |
| Chemical compound, drug | L-Lysine:2HCL (13C6, 99%; 15N2, 99%) | Cambridge Isotope Laboratories | CNLM-291-H-0.25 | MS assays |
| Chemical compound, drug | Iodoacetamide | Sigma-Aldrich | I1149 | MS assays and protein purification |
| Chemical compound, drug | Ammonium bicarbonate | Sigma-Aldrich | 9832 | MS assays |
| Chemical compound, drug | Formic acid 99–100% | VWR Chemicals | 20318.297 | MS assays |
| Chemical compound, drug | iRT Kit | Biognosis | Ki-3002-1 | MS assays |
| Chemical compound, drug | BioPureSPN MINI Columns Silica C18 | The Nest Group, Inc | HUM S18V | MS assays |
| Chemical compound, drug | BioPureSPN MACRO Columns Silica C18 | The Nest Group, Inc | HMM S18V | MS assays |
| Antibody | α-V5 (mouse monoclonal) | Invitrogen | R960-25 | Western blotting, 1:2000 |
| Antibody | α-Xpol (serum, polyclonal) | In house facility | In house facility | Western blotting, 1:5000 |
| Antibody | α-GFP (mouse monoclonal) | Roche | 1181446000 | Western blotting, 1:2000 |
| Antibody | α-Hexokinase (rabbit monoclonal) | US Biologicals | H2035-01 | Western blotting, 1:3000 |
| Antibody | α-Mouse IgG Alexa Fluor 680 (goat polyclonal) | Thermo Fisher Scientific | A-21057 | Western blotting, 1:10,000 |
| Antibody | α-Rabbit IgG IRDye800CW (goat polyclonal) | Li-COR Biosciences | 926-32211 | Western blotting, 1:10,000 |
| Antibody | anti-PGK1 (22C5D8) | Abcam | ab113687 | Western blotting, 1:5000 |
| Antibody | anti-HA | Biolegend | # MMS-101P | Western blotting, 1:5000 |
| Software, algorithm | NuRim | Rajoo et al., 2018; Vallotton et al., 2019 | / | Data analysis |
| Software, algorithm | ImageJ/Fiji | (Schindelin et al., 2012) | / | Data acquisition |
| Software, algorithm | NIS Elements | Nikon | / | Data acquisition |
| Software, algorithm | MATLAB | MathWorks | / | Data analysis |
| Software, algorithm | AlphaFold2.1.1 | (Jumper et al., 2021) | / | Data analysis |
| Software, algorithm | AlphaFold visualization | https://colab.research.google.com/drive/1CizC7zmYvFkav5qfBbWxhgUHRoxwym2w | / | Data analysis |
| Software, algorithm | Excel | Microsoft | / | Data analysis |
| Software, algorithm | R v. 4.1.2 | R Project | / | Data analysis |
| Software, algorithm | Prism 7, Prism 9 | GraphPad | / | Visualization |
| Software, algorithm | Illustrator v. 26.0.3 | Adobe | / | Visualization |

### Yeast culturing conditions

Yeast strains were grown to mid-log phase (OD_600_ 0.4–0.9) for at least 12 hours at 30°C. For proteomic analysis, western blotting, and immunogold electron microscopy, cells were cultured in rich YPD medium (1% yeast extract, 2% peptone, 2% dextrose). For fluorescence microscopy, growth curves, and spotting assays, cells were grown in synthetic complete medium (SCD; 6.7 g/L yeast nitrogen base without amino acids, 2% dextrose) supplemented with the appropriate amino acids and nucleobases.

For acute depletion using the auxin-inducible degron (AID) system, log-phase cultures of strains expressing Jjj2-AID or Brl1-AID were treated with 4 μM IP6 (phytic acid dipotassium salt; Sigma-Aldrich, P5681) and 500 µM auxin (indole-3-acetic acid dissolved in ethanol; Sigma-Aldrich, I2886) for the durations specified in the respective figure legends.

Strains containing galactose-inducible constructs driven by the GAL1 promoter were pre-cultured in SC medium supplemented with 2% raffinose. Expression was subsequently induced by adding 2% galactose to log-phase cultures. As an exception, for *in cellulo* experiments utilizing the GFP-Nup100FG construct, expression was induced with 1% (w/v) galactose for 2 hours and then repressed with 2% (w/v) glucose for 4 hours prior to imaging. Immediately before imaging, cells were incubated for 2 minutes with Calcofluor (Fluorescent Brightener 28; 5 µg/mL final concentration) to stain the cell wall and facilitate cell recognition.

Strains expressing NUP82-RITE(Red-to-Green) constructs were grown to mid-log phase in synthetic medium lacking uracil (SD -Ura) to select for non-recombined cells. To induce Cre-mediated recombination prior to imaging, cultures were supplemented with 1 µM β-estradiol (Sigma-Aldrich, E8875) and 20 mg/mL uracil.

Recombination was induced 30 minutes prior to acute depletion of Jjj2-AID with auxin to ensure that the genomic switch was complete before Jjj2 levels decreased and condensates began to form. Cells were imaged 5 hours and 30 minutes after the switch, a time point when both “old” (red) and “new” (green) Nup82 populations were sufficiently enriched at the NE to form a distinct rim. This clear demarcation of the nuclear boundary was essential to unequivocally distinguish cytoplasmic condensates from the NE NPC signal.

For the 1,6-hexanediol sensitivity assays, log-phase cells were incubated in SCD medium containing 5% 1,6-hexanediol (Sigma-Aldrich, 240117) for 10 minutes immediately prior to imaging.

### Affinity pulldown coupled to mass spectrometry

#### · Cell lysis and affinity purification

All steps were performed on ice. Frozen yeast pellets were resuspended in 1 mL lysis buffer (20 mM HEPES pH 7.5, 50 mM KOAc, 20 mM NaCl, 2 mM MgCl2, 1 mM DTT, 10% v/v glycerol) in 2 mL screw-cap micro tubes (Sarstedt) containing ∼1 mL of 0.5 mm glass beads (BioSpec). Tubes were filled completely with lysis buffer, avoiding air bubbles. Cells were lysed using a mini BeadBeater-24 (BioSpec) for four 1-min cycles at 3500 oscillations/min, with 1-min cooling intervals in ice water. Lysates were cleared by centrifugation (15’000 x g, 30 s). For affinity purifications, 1 mL of supernatant was mixed with 110 µL of 10× detergent mix (protease inhibitor cocktail (Sigma-Aldrich), 5% v/v Triton X-100, 1% v/v Tween-20 in lysis buffer) and 1 mg IgG Dynabeads (pre-equilibrated in lysis buffer containing 0.5% Triton X-100 and 0.1% Tween-20). Following a 30-min incubation with constant agitation at 4°C, beads were washed twice with 1 mL wash buffer (0.1% v/v Tween-20 in lysis buffer). Proteins were eluted in 40 µL 1× Laemmli buffer (2 min, 50°C), completely denatured (5 min, 95°C), and snap frozen.

#### · In-gel tryptic digestion

Elutes were concentrated in a 4% SDS-PAGE stacking gel, stained with SimplyBlue SafeStain (Invitrogen, LC6060), and gels were washed in distilled water for 1 h. Excised protein bands were reduced with 6.5 mM DTT in 100 mM ammonium bicarbonate (1 h, 60°C), alkylated with 54 mM iodoacetamide in 100 mM ammonium bicarbonate (30 min, 30°C, in the dark), and digested with 1.25 µg sequencing-grade porcine trypsin (Promega) in 100 mM ammonium bicarbonate (overnight at 37°C). Peptides were desalted using C18 BioPureSPN mini columns (The Nest Group), washed thrice with Buffer A (0.1% formic acid in HPLC-grade water), and eluted thrice with 150 µL Buffer B (50% acetonitrile, 0.1% formic acid). Peptides were recovered in 12.5 µL Buffer A containing iRT peptides (1:50 v:v, Biognosys).

#### · Mass spectrometry acquisition

Peptides were analyzed using an Orbitrap Exploris 480 mass spectrometer (Thermo Scientific) coupled to a Vanquish Neo UHPLC system (Thermo Scientific). Separation was performed with a C18 reversed phase column (75 μm x 400 mm (New Objective), packed in-house with ReproSil Gold 120 C18, 1.9 μm (Dr. Maisch GmbH)), using a 120-minute linear gradient of 7-35% Buffer B (0.1% formic acid, 80% acetonitrile) at a flow rate of 300 nL/min. The mass spectrometer was operated in data-independent acquisition (DIA) mode. Full MS1 scans were acquired from 350-1150 m/z at 120’000 resolution (250% normalized AGC target, 264 ms maximum injection time). This was followed by 41 variable MS2 windows (350-1150 m/z,1 m/z overlap) at 30000 resolution (250% normalized AGC target, 64 ms maximum injection time). Fragmentation was performed via higher-energy collisional dissociation (HCD) with a normalized collision energy (NCE) of 28%.

#### · DIA MS data analysis

DIA (Data-independent acquisition) data were analyzed using Spectronaut v15 (Biognosys AG) via the directDIA workflow. Spectra were searched against the Saccharomyces cerevisiae protein database (downloaded from the Saccharomyces Genome Database on 13/10/2015; 6,713 entries). Search parameters included carbamidomethylation of cysteine as a fixed modification, while methionine oxidation and N-terminal acetylation were included as variable modifications. Only tryptic peptides with a maximum of two missed cleavages were considered. For data extraction, default Spectronaut settings were applied, except for disabling “cross-run normalization”. Fragment-level ion intensities were exported and processed in R. Low-quality fragments were excluded based on Spectronaut’s “F. ExcludedFromQuantification” flag. Precursor intensities were calculated as the sum of fragment ion intensities, and only prototypic precursors were retained. To ensure robust quantification, precursors were further filtered to include only those present across all three biological replicates, and the top three most intense precursors were summed to determine protein intensities. For proteins not detected in the mock ZZ pull-downs, missing values were imputed by sampling from a normal distribution fitted to the bottom 5% of the ZZ intensity range. Samples were median-normalized, and final protein intensities were calculated as the mean of three biological replicates. Statistical significance was assessed using a two-tailed Student’s t-test and resulting p-values were adjusted for multiple testing using the Benjamini–Hochberg correction.

To identify proteins specifically associated with assembling NPCs, a previously published affinity purification-mass spectrometry dataset comprising ten different Nup baits was utilized (Onischenko et al., 2020, Cell). The differential enrichment of co-purified proteins was compared between early-assembling baits (Nup188, Nup57, Nup84, Nup133) and a late-assembling bait (Mlp1). For each individual bait, protein intensities were first summarized as the median of three biological replicates. Next, a consensus intensity for each assembly tier was determined by taking the median of all baits within that tier, and the fold change between the early and late assembly tiers was calculated. Finally, Nups and NTRs were excluded from the dataset to focus the analysis exclusively on non-NPC interactors.

### Western blotting

For each condition, 5mL of yeast culture in mid-log phase was collected by centrifugation and lysed by a 15 min incubation in 0.1 M sodium hydroxide at room temperature. Then cells were pelleted, resuspended in 50 µL Laemmli sample buffer (62.5 mM Tris–HCl pH 6.8, 10% glycerol, 2% SDS, 5% 2-mercaptoethanol, 100 mM DTT, and 0.04% bromophenol blue) and heat denatured for 5 min at 95°C. An equivalent of 1 OD_600_ unit of cellular material was loaded per well. Proteins were electrophoretically separated using either custom-made 10% polyacrylamide gels or commercial 4%–12% gradient gels (NuPAGE 4–12% Bis-Tris, 1.0 mm; Thermo Fisher Scientific, NP0321). Proteins were subsequently transferred in 1x TBE buffer to a nitrocellulose membrane (Amersham Protran 0.2 NC; Cytiva, 10600015) using a Trans-Blot Turbo semi-dry transfer system (Bio-Rad). Prior to antibody incubation, membranes were blocked for 1 h in 5% PBST-milk (1× PBS, 0.1% Tween-20, 5% dry milk). Then, membranes were incubated with primary antibody for 1 hr at room temperature or overnight at 4°C, washed three times for 10 min in PBST (1× PBS pH 7.4; 0.1% Tween-20) followed by a 1 h incubation with secondary antibody at room temperature. Membranes were washed again three times for 10 min in PBST before fluorescence signal was imaged with the CLx ODYSSEY (Li-COR). Primary antibodies used were: mouse monoclonal α-GFP (Roche, 11814460001, 1:2000), mouse monoclonal α-V5 (Invitrogen, R960-25; 1:2000), serum polyclonal α-XpoI (in house facility), rabbit monoclonal α-hexokinase (US Biologicals, H2035-01; 1:3000), mouse monoclonal α-PgkI (Abcam, ab113687), mouse monoclonal anti-HA (Biolegend, # MMS-101P, 1:5000). Secondary antibodies used were goat α-mouse IgG Alexa Fluor 680 (Thermo Fisher Scientific, A-21057; 1:10,000) and goat α-rabbit IgG IRDye800CW (Li-COR Biosciences, 926-32211; 1:10,000).

For western blot quantification, band intensity measurements were performed using ImageJ software (Schindelin et al., 2012). The signal intensity of the target protein was normalized to that of the respective loading control. For all measurements, identical regions of interest were applied across lanes, and background values were subtracted.

### Spotting assay

For growth phenotypic analyses of the J-domain protein deletion mutants, yeast strains were grown to saturation in YPD medium. For strains harboring galactose-inducible *JJJ2* derivatives, cells were grown to log phase in SC medium supplemented with 2% raffinose.

In all cases, cultures were adjusted to an initial OD_600_ of 0.2, from which fivefold serial dilution series were prepared. For each dilution, 4 uL of culture was spotted onto the appropriate agar plates: YPD for the deletion mutants, and synthetic media supplemented with either 2% galactose (induction) or 2% glucose (repression) for the overexpression strains. Plates were incubated at 30°C until adequate colony growth was observed.

### Growth curves

Starting from cell cultures grown in SCD to log phase, cultures with OD_600_=0.15 were prepared in NUNC 24-well plates. Growth curves were recorded using a CLARIOstar® multimode microplate reader (BMG Labtech, Germany) in absorbance mode. Absorbance measurements at 600nm were taken every 5 minutes at 30°C. Each well was read using 20 flashes per cycle, with orbital averaging enabled and a scan diameter of 8 mm. Prior to each cycle, plates were shaken with double orbital movement at 500 rpm for 60 seconds. Additional shaking between readings was also included.

### Fluorescence microscopy of yeast cells

For live-cell fluorescence imaging, yeast cells were immobilized in 384-well glass-bottom plates (MatriPlate; Brooks Life Sciences) pre-coated with concanavalin A (Sigma-Aldrich). Imaging was performed at 30°C using a x100 Plan-Apo VC objective (NA 1.4; Nikon) on a Nikon Eclipse Ti inverted epifluorescence microscope. The system was equipped with a Spectra X LED light source (Lumencor) and controlled via NIS-Elements software (Nikon). Images were captured using a Flash 4.0 sCMOS camera (Hamamatsu) and subsequently processed and analyzed using ImageJ software.

For GFP-Nup100FG experiments in cellulo, cells were imaged at 30°C using a DeltaVision Elite deconvolution microscopy system (Applied Precision) equipped with a 100× Olympus UPLS Apo oil-immersion objective (NA 1.4) and a CoolSNAP HQ2 CCD camera. Light excitation was provided by an InsightSSI solid-state illumination module, utilizing FITC (525/48 nm), A594 (625/45 nm), and DAPI (435/48 nm) filter sets for channel selection. To capture the full volume of the cells, three-dimensional fluorescence Z-stacks were acquired across 30 focal planes with a 0.2-μm step size. A single brightfield reference image was also recorded at the mid-plane of the sample using polarized light.

### Quantitative image analysis of fluorescence microscopy images of yeast cells

#### · Quantification of Nup-GFP intensity at the nuclear envelope

For comparisons of Nup-GFP intensity at the NE between wild-type (WT) and jjj2Δ (Fig. 1E), the automated image analysis pipeline NuRim was utilized to quantify the fluorescence intensity signal at the NE for various Nup-GFP fusion proteins (Rajoo et al., 2018; Vallotton et al., 2019). Briefly, nuclear contours were defined in an unbiased manner based on the fiducial marker Hmg1-mCherry. The fluorescence intensities of NUP-yEGFP along these contours were subsequently extracted using ImageJ and plotted in GraphPad Prism.

For experiments comparing Nup-yEGFP intensity at the NE during Jjj2 overexpression (Fig. S9, B), the automated NuRim pipeline could not be used due to increased vacuolar autofluorescence, which caused segmentation artifacts along the NE contour. Therefore, these quantifications were performed manually. Nuclear contours were traced using the segmented line tool in ImageJ (spline fit enabled, line width: 3). For each contour, the mean intensity was measured and corrected by subtracting the mean cytoplasmic intensity obtained from a manually drawn reference line. For each biological replicate, 50 nuclear contours were quantified per strain and condition, and the average values were calculated and plotted in GraphPad Prism.

#### · Cytoplasmic to nuclear intensity ratio quantification

To quantify cytoplasmic-to-nuclear fluorescence intensity ratios (Fig. 5D; Fig. S7E, H, K), cellular and nuclear segmentation was performed using the deep learning-based tool Cellpose (version 3.1.1.1). Cell segmentation from brightfield images was performed using the pretrained *cyto2* model with the following parameters: diameter = 75, minimum object size = 3, verbose output enabled, exclusion of edge objects, no NPY output, cell probability threshold = 1, and flow threshold = 2. Nuclear segmentation from the Nup82-mCherry fluorescence channel employed the same *cyto2* model with adjusted parameters: diameter = 20, minimum object size = 3, cell probability threshold = 3.5, and flow threshold = 0.

Fluorescence intensities were subsequently quantified using CellProfiler 4. Cell and nuclear masks generated by Cellpose were imported to define primary ‘Cell’ and ‘Nucleus’ objects. Cells were filtered based on area (1000–10000), form factor (0.7–1), and mean intensity (0.1–1). Nuclei were filtered by area (100–1000) and form factor (0.7–1). To exclude peripheral regions, filtered cell masks were shrunk by 4 pixels, and cytoplasmic regions were defined by subtracting the nucleus from the shrunken cell mask. To further exclude regions near the plasma membrane and NE, both cytoplasmic and nuclear masks were shrunk by an additional 3 pixels. A single manually drawn region per image was used to define background fluorescence. Mean intensities were measured in the shrunken cytoplasmic, nuclear, and background areas. Background-subtracted intensities from cytoplasm and nucleus were used to compute the cytoplasmic-to-nuclear ratio. Only cells with both nuclear and cytoplasmic segmentations were included in the analysis.

#### · Pearson’s correlation analysis

For co-localization quantifications (Fig. 2E-F), NE contours were manually delineated based on the Nup-mCherry signal, and intensity profiles were extracted using Fiji. The Pearson’s correlation coefficient between intensity values in the green and red channels was calculated along these contours. For each biological replicate, 50 nuclear contours were analyzed per strain and condition.

#### · Quantification of the percentage of cells with Nup-GFP cytoplasmic foci

The percentage of cells displaying cytoplasmic Nup-GFP foci was determined by manual counting in a blinded manner.

#### · Image analysis of Nup100FG and TDP-43-GFP condensation assays

For the analysis of Nup100FG condensates in Fig. 4A-G and TDP-43-GFP condensates in Fig. S5D-E, the PhaseMetrics plugin was used, as described in (Bergsma et al, 2025). PhaseMetrics performs automatic detection and analysis of several particle properties, including particle intensity, size and circularity. A maximum intensity Z-projection is used for creating a segmented image for object detection, after which the object masks are re-directed to the sum of slices projection to extract the desired measurements.

GFP-Nup100FG expressing cells were classified based on the GFP signal as Soluble/NS (no particles), Condensates (1, 2, or ≥3 round particles), or Aggregates (non-round/irregular particles). Cells containing both round and non-round particles were classified as Aggregates. Particle classification criteria were:

|  | Circularity |  | Perimeter (μm) |  | Area (μm<sup>2</sup>) |  |
| --- | --- | --- | --- | --- | --- | --- |
| Category | Min | Max | Min | Max | Min | Max |
| Aggregates | 0 | 0.6 | 2.5 | 1000 | 0.5 | 1000 |
| Condensates | All particles that do not meet the above criteria |  |  |  |  |  |

Because TDP-43-GFP does not form elongated condensates, the aggregate category was omitted from that specific analysis. Particle classification criteria were:

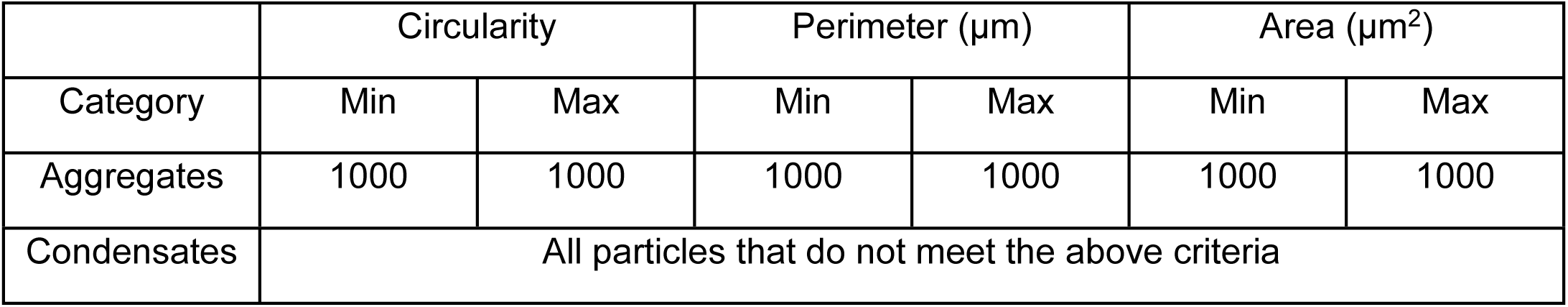

|  | Circularity |  | Perimeter (μm) |  | Area (μm<sup>2</sup>) |  |
| --- | --- | --- | --- | --- | --- | --- |
| Category | Min | Max | Min | Max | Min | Max |
| Aggregates | 1000 | 1000 | 1000 | 1000 | 1000 | 1000 |

|  |  |
| --- | --- |
| Condensates | All particles that do not meet the above criteria |

### Immunogold assay and electron microscopy (EM)

For immunogold-EM experiments (Fig. 2G), sample preparation and labeling were performed largely as described previously (Panagaki et al., 2021). For morphology-only EM experiments (Fig. S2F and Fig. S9C), the same protocol was followed, omitting the immunolabeling steps.

Yeast cells were cultured in SCD medium at 30°C. Degradation of Brl1 in log-phase yeast cultures was induced by the addition of 4 μM IP6 and 500 µM auxin for approximately 5 h. The subcellular localization of endogenous Jjj2, C-terminally tagged with yEGFP, was subsequently visualized via immunogold electron microscopy.

#### · High-pressure freezing and freeze substitution

Yeast cells were high-pressure frozen (Wohlwend HPF Compact 3, Sennwald, Switzerland) followed by freeze substitution in 2% uranyl acetate (UA) dissolved in acetone and 10% methanol, and embedded into HM20 lowicryl resin (Polysciences) that was UV polymerized at -50°C. The polymerized resin blocks were sectioned at a thickness of 70 nm using an ultramicrotome, and the resulting sections were collected on 3 mm copper grids (200 mesh) coated with 1% Formvar (vinylec E).

#### · Immunolabelling for electron microscopy

For immunogold labeling, grids carrying 70 nm sections were fixed in 1% paraformaldehyde in PBS for 10 minutes and subsequently blocked for 1 hour with a solution containing 0.1% fish skin gelatin and 0.8% BSA in PBS. Sections were then incubated with a primary α-GFP antibody (diluted 1:5 in blocking buffer) overnight at 4°C.

Following the primary incubation, grids were washed three times for 20 minutes each in PBS, incubated for 1 hour with a gold-conjugated secondary antibody (diluted 1:20 in blocking buffer), and washed again three times for 20 minutes each in PBS. The labeled sections were post-fixed in 2.5% glutaraldehyde for 1 hour, washed three times for 1 minute each in distilled water, and finally contrast-stained using 2% UA and Reynolds’ lead citrate (Reynolds et al., 1963).

#### · Electron microscopy image acquisition

All thin sections were imaged at 120 kV on a Tecnai 12 (Thermofisher) equipped with a 4k x 4k Ceta camera and a Lab6 electron source.

#### · Immunolabelled electron microscopy image quantification

To quantify Jjj2-GFP labeling density (Fig. 2H), the surface area and total number of gold particles within distinct cellular compartments were measured. Defined compartments, including the entire cell, nucleus, NE, lipid droplets, and herniations, were manually delineated using ImageJ software. To account for the physical dimensions of the primary-secondary antibody-gold particle complex (approximately 30 nm), each defined compartment boundary was expanded by 30 nm in all directions. Because membrane curvature at defective NPC assembly sites (herniations) was not always distinct, these structures were not delineated by the membrane bilayer itself; instead, highly electron-dense regions localized along the nuclear envelope (which were absent in the non-degraded Brl1 control conditions) were classified as herniations.

### Protein purification and *in vitro* assays

#### · Protein purification

To prepare the proteins analyzed in Figure S6, we expressed and purified the yeast Nup100 FG-domain along with the molecular chaperone Jjj2 according to previously described protocols (Kuiper et al., 2022). Briefly, the Nup100FG fragment (aa 1–580), full-length Jjj2 (FL), and the truncated Jjj2(aa 1–132) variant were expressed in *E. coli* using the pSF350 plasmid, which encodes an N-terminal His6-tag and a C-terminal cysteine residue.

Recombinant expression was carried out in 200 mL *E. coli* cultures grown at 37°C until reaching an OD600 of 0.5-0.8. At this point, cultures were induced with 0.5 mM isopropyl-β-D-thiogalactoside (IPTG) for 5 hours at 20°C. Following induction, cells were harvested via centrifugation (4,500g, 15 min, 4°C), and the resulting pellets were preserved at -80°C.

For purification, frozen bacterial pellets were thawed and resuspended in 10 mL denaturing lysis buffer (100 mM Tris-HCl and 2 M guanidine-HCl, pH 8.0; Thermo Scientific, #24110) supplemented with 1 mM phenylmethylsulfonyl fluoride (Serva, #32395.02), 5 mM DL-1,4-dithiothreitol (DTT; 99%, ACROS Organics, #10215550), and a protease inhibitor cocktail (cOmplete ULTRA Tablets, Mini, EDTA-free, EASYpack Protease Inhibitor Cocktail, #5892791001). Mechanical disruption of cells was executed with 0.1 mm glass disruptor beads (Scientific Industries, #SI-BG01) using a FastPrep instrument. To clarify the lysates, we performed two successive centrifugation steps at 20,000g: an initial 5-minute spin followed by a 15-minute spin of the recovered supernatant. This cleared supernatant was equilibrated in lysis buffer and incubated with Ni Sepharose 6 Fast Flow beads (Cytiva, #17-5318-03) for 1 hour at 4°C. The slurry was transferred to Poly-Prep Chromatography Columns (Bio-Rad, #7311550), and the matrix was washed twice with 10 mL of lysis buffer (100 mM Tris-HCl and 2 M guanidine-HCl, pH 8.0) containing 2.5 mM DTT and 50 mM imidazole (PUFFERAN ≥99%, Carl Roth, #X998.4). Target proteins were then eluted in elution buffer containing 500 mM imidazole (100 mM Tris-HCl, 2 M guanidine-HCl, pH 8.0, and 10% glycerol).

For fluorophore-labeling experiments, the purification process was modified slightly to remove residual reducing agents. Specifically, the second column wash was performed with 10 mL of DTT-free lysis buffer (100 mM Tris-HCl, 2 M guanidine-HCl, pH 8.0) containing 50 mM imidazole (PUFFERAN ≥99%, Carl Roth, #X998.4). Following washing, the immobilized proteins were incubated on the Ni-sepharose matrix for 1 hour at room temperature with 4.5 µg of either fluorescein-5-maleimide (Thermo Scientific, #62245) for the FG-Nups, or Alexa Fluor 594 C5 Maleimide (Thermo Fisher, #A10256) for the Jjj2 variants. Unbound dyes were washed away with lysis buffer, and the labeled proteins were eluted in a buffer comprising 500 mM imidazole, 100 mM Tris-HCl, 2 M guanidine-HCl (pH 8.0), and 10% glycerol.

To prevent spontaneous disulfide bond formation, the sulfhydryl groups of the purified proteins were reduced using 10 mM DTT for 30 minutes at 50°C and subsequently blocked with 15 mM iodoacetamide alkylating reagent (Sigma-Aldrich, #I1149-25G) for 30 minutes at room temperature. For sample concentration, the FG-Nup fragments were concentrated in Vivaspin Protein Concentrator spin columns (Vivaspin 2, 10/30 kDa MWCO Polyethersulfone, Cytiva, #28-9322-47, #28-9322-48). In contrast, the full-length Jjj2 and Jjj2(aa 1–132) constructs were concentrated using Amicon Ultra-4 Centrifugal Filter Units (Amicon Ultra-4, 3 and 30 kDa MWCO regenerated cellulose, MilliporeSigma, #UFC800324, #UFC803024) following the manufacturers’ protocols. Final protein yields were quantified using the Pierce BCA Protein Assay Kit (Thermo Scientific, #23227) in accordance with the manufacturer’s directions, utilizing Pierce Bovine Serum Albumin Standard Ampules (2 mg mL⁻¹, #23209) to generate the standard curve.

#### · Sample preparation for *in vitro* phase separation assays

In Fig. S6C-F, the phase separation behavior of purified proteins was assayed. The purified proteins were diluted to a protein concentration of 3 μM in assay buffer in low-protein-binding tubes for one hour at room temperature. The assay buffer comprises 50 mM Tris-HCl, 150mM NaCl, pH8.0, supplemented with 10% w/v of polyethylene glycol 3350 (PEG3350; Sigma-Aldrich, P4338-2 KG).

To minimize the impact of the fluorescent labels, labeled and unlabeled batches of purified proteins were mixed prior to the experiments. Jjj2 full-length and Jjj2(aa1–132) were mixed at a 1:20 ratio, meaning that maximally 5% of the molecules were labeled, whereas Nup100FG was mixed at a 1:5 ratio, resulting in a maximum of 20% labeled molecules.

To assess the effect of Jjj2 full length and Jjj2(aa1-132) on FG-Nup particle properties, 3 μM of 5MF labeled Nup100FG and AF594 labeled Jjj2 full length or Jjj2(aa1-132) were mixed at a 1:1 ratio and left to phase separate for 1 h at room temperature, prior to microscopy assessment.

#### · Fluorescence microscopy for *in vitro* phase separation assays

For imaging of in vitro purified proteins in Fig. S6C-F, 2 μL samples of phase separated protein mixtures were mounted on untreated glass slides with coverslip. Microscopy was performed in a temperature-controlled environment at 20°C using a DeltaVision Deconvolution Microscope (Cytiva), using an Olympus UPLS Apo 100× oil-immersion objective (NA 1.4) and InsightSSITM Solid State Illumination using the FITC 525/48 and A594 625/45 filter sets. Detection was done with a CoolSNAP HQ2 or EDGE sCMOS5.5 camera. Fluorescence images were acquired with 30 Z-slices of 0.2 μm. Images were deconvolved using softWoRx software (Cytiva) and processed using ImageJ.

#### · Image analysis of *in vitro* phase separation assays

For the analysis of particle properties in Fig. S6E-F, the PhaseMetrics plugin was used, as described in (Bergsma et al., 2025).

#### · Filter trap assays

For filter trap assays in Fig. S6A-B, protein samples were prepared as explained above, after which 180 μL of sample buffer supplemented with 0.5% SDS was added to the sample and mixed by vortex before loading onto the Bio-Dot apparatus (Bio-Rad). FTAs were performed as described before (Kuiper et al, 2022). After this, membranes were blocked with 2.5% BSA in PBS-T (0.1%), incubated overnight with mouse primary antibody anti-His (1:5000, monoclonal mouse Tetra-His antibody, Qiagen, #34670), washed three times with PBS-T, incubated for 1h with secondary anti-mouse m-IgGk BP-HRP protein (Santa Cruz, sc-516102) (1:2500) and washed three times with PBS-T. Chemiluminescence was detected using enhanced chemical luminescence (ECL) reagent using the Chemidoc imaging system (BioRad). For the quantification of the FTA, band intensities were measured using FIJI and expressed relative to the average intensity of the control.

### Western blot and filter trap assays with GFP-Nup100FG construct in yeast cells

For filter trap assays in Fig. 4D-E, cells were washed once in cold PBS and scraped in 200 μL lysis buffer (25 mM HEPES, 100 mM NaCl, 1 mM MgCl_2_, 1% NP-40 (Igepal CA-630, Sigma), EDTA-free complete protease inhibitors cocktail (Roche) and Benzonase (EMD Millipore) (∼90 units mL^−1^)), and left on ice for minimum of 30 min with intermittent vortexing until chromatin was dissolved. Protein concentrations were determined using DC protein assay (Bio-Rad). Concentrations were equalized and diluted in filter trap assay buffer (10 mM Tris–HCl pH 8.0, 150 mM NaCl and 50 mM dithiothreitol, 0.5% SDS), boiled for 5 min and prepared in three five-fold serial dilutions into a final of 1×, 5× and 25× diluted samples for FTA. FTA samples were loaded onto a 0.2-μm-pore-size cellulose acetate membrane pre-washed with 0.1% SDS-containing FTA buffer. Membranes were washed two times with 0.1% SDS-containing FTA buffer, blocked with 5% non-fat milk, and blotted with anti-GFP/YFP (Clontech, JL-8). After HRP-conjugated secondary antibody incubation, visualization was performed using enhanced chemiluminescence and ChemiDoc Imaging System (Bio-Rad). Bands were measured (ImageLab 5.2.1), and the measurement of the three dilutions was averaged. Values normalized to control were plotted in a graph using GraphPad Prism.

## SUPPLEMENTARY FIGURES

**Figure S1:**
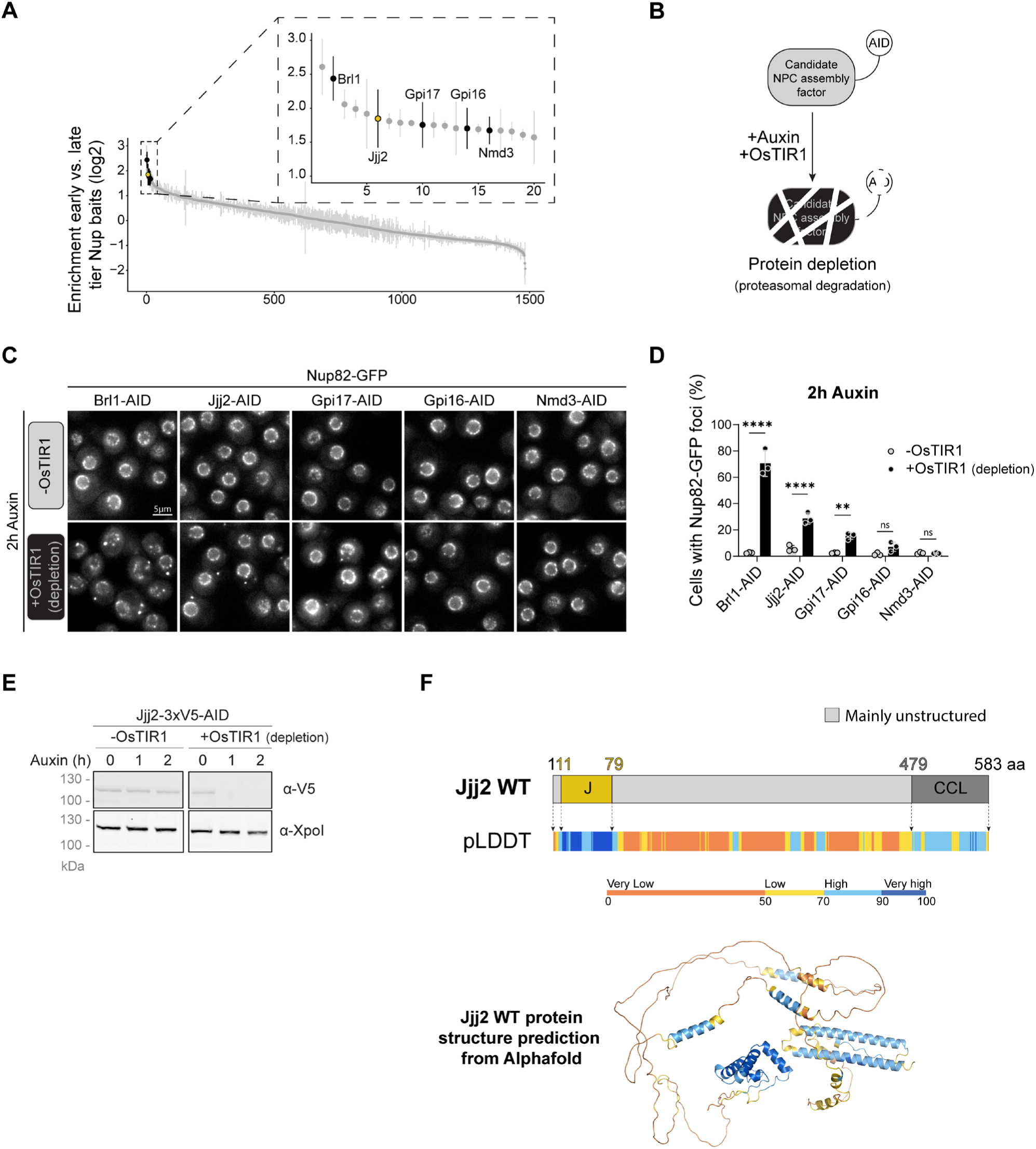
Jjj2 is identified as a putative Nup regulator. **(A)** Identification of putative Nup regulators. Ranking of 1,500 proteins co-purified with 10 nucleoporin baits (Onishchenko et al., 2020, Cell) based on fold enrichment differences between early and late assembling baits. Inset shows the top 20 early-enriched proteins; targets selected for fluorescence validation are highlighted in black. Data represent median ± SD of three biological replicates. **(B)** Schematic of the Auxin-Inducible Degron (AID) system. Putative assembly factors were C-terminally tagged with an IAA7 motif (AID) in strains expressing the auxin-sensitive F-box protein OsTIR1 (+OsTIR1) or lacking it (-OsTIR1; control). Auxin addition triggers the recruitment of the SCF-TIR1 E3 ubiquitin ligase complex, leading to the proteasomal degradation of the targeted protein. **(C)** Representative fluorescence micrographs of Nup82-GFP in control (-OsTIR1) and putative Nup regulator depleted (+OsTIR1) cells. Scale bar: 5 μm. **(D)** Percentage of cells in (C) showing cytoplasmic Nup82-GFP foci. Data represent mean ±SD of three biological replicates. Statistical test: Two-way ANOVA with Sidak’s multiple comparison test. **p < 0.01, ****p < 0.0001. **(E)** Western blot monitoring the degradation of Jjj2-3xV5-IAA7 (Jjj2-AID) at the indicated time points following auxin addition. Jjj2 was detected using an anti-V5 antibody; anti-Xpo1 served as a loading control. **(F)** Structural organization of Jjj2. Top: Schematic representation of the Jjj2 domain architecture, with amino acid positions indicated. The J-domain (J) is highlighted in yellow, the coiled-coil-like (CCL) domain in dark grey and predicted intrinsically disordered regions (areas with low Predicted Local Distance Difference Test, or pLDDT, scores in AlphaFold 2) in light grey. The lower panel displays the corresponding pLDDT score distribution of the AlphaFold 2 model plotted across the primary sequence. Bottom: ribbon diagram of the AlphaFold2 structural prediction for Jjj2, with residues color-coded according to their respective pLDDT scores.

**Figure S2:**
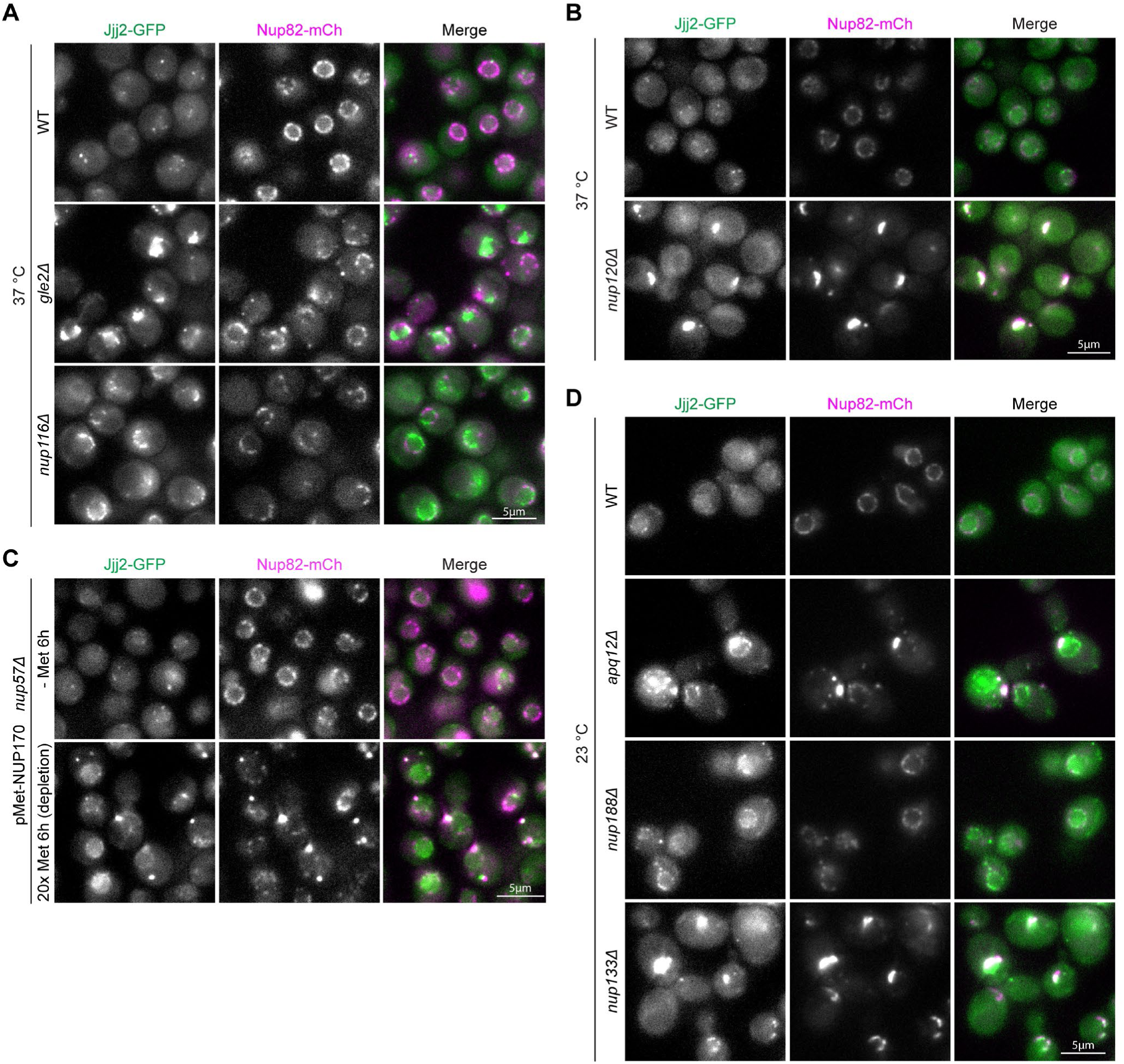
Jjj2 localizes to defective NPCs. (A-B) Representative fluorescence micrographs of WT, *gle2Δ*, *nup116Δ* and *nup120Δ* cells expressing Jjj2-GFP and Nup82-mCherry. Cells were grown at 23 °C and shifted to 37 °C for 3 h prior to imaging. Scale bar: 5 μm. **(C)** Representative fluorescence micrographs of pMet-NUP170 *nup57Δ* cells expressing Jjj2-GFP and Nup82-mCherry. Cells were initially cultured in medium lacking methionine (-Met) and shifted to medium supplemented with 20x methionine (20x Met) to repress NUP170 transcription for 6 h prior to imaging. Scale bar: 5 μm. **(D)** Representative fluorescence micrographs of WT, *apq12Δ*, *nup188Δ* and *nup133Δ* cells expressing Jjj2-GFP and Nup82-mCherry and grown at 23 °C. Scale bar: 5 μm.

**Figure S3:**
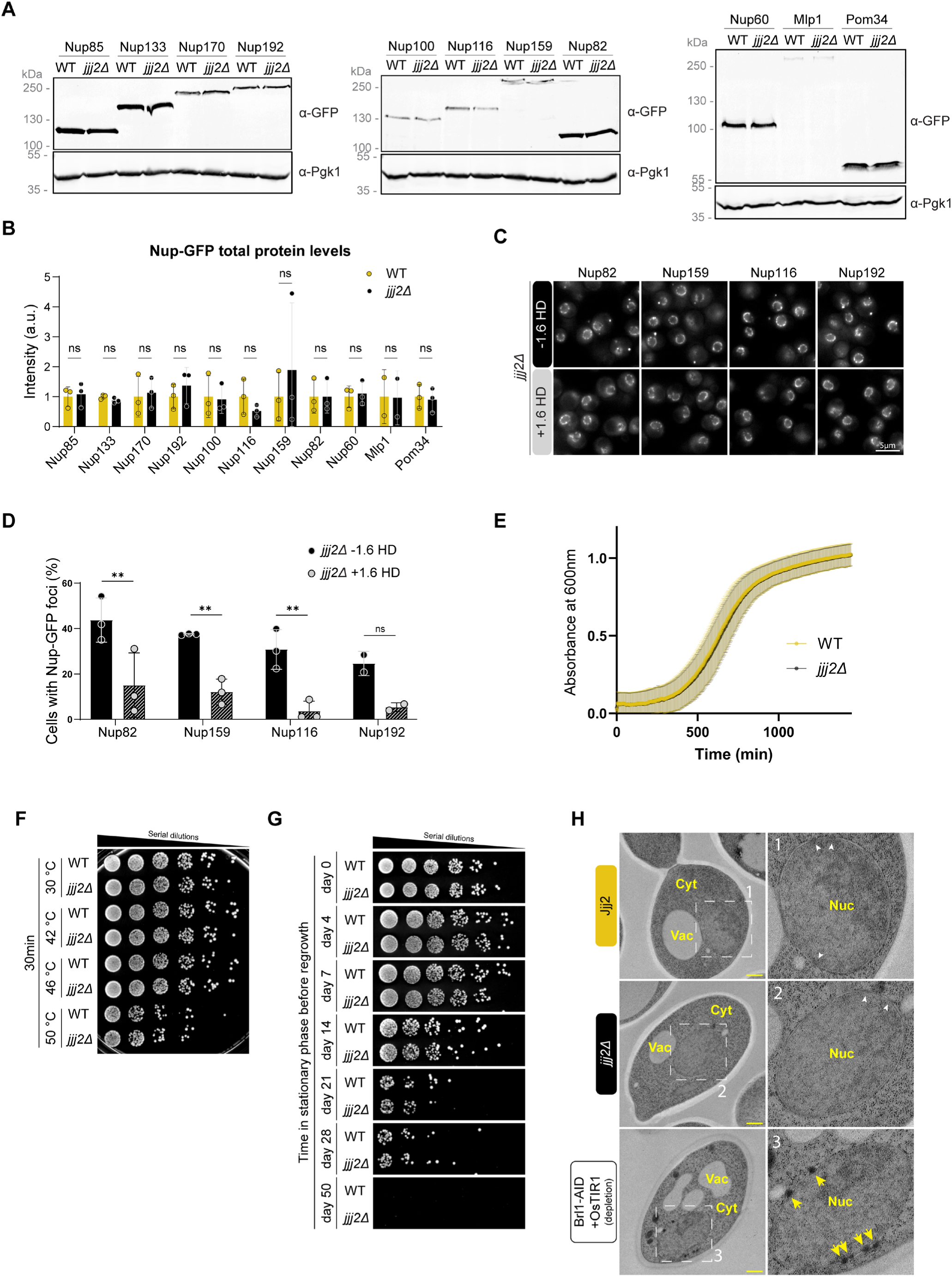
Cells lacking Jjj2 exhibit normal growth, baseline nucleoporin levels and no obvious NPC abnormalities. **(A)** Representative western blots showing levels of the indicated GFP-tagged Nups in whole-cell protein extracts from WT and *jjj2Δ* cells. Nups were detected using an anti-GFP antibody; anti-Pgk1 served as a loading control. **(B)** Band intensities from (A) were normalized to the Pgk1 loading control. Data represent mean ±SD from three biological replicates. Statistical test: Two-way ANOVA with Sidak’s multiple comparison test. ns = non-significant. **(C)** Representative fluorescence micrographs of GFP-tagged Nups in *jjj2Δ* cells following treatment with or without 5% 1,6-hexanediol for 10 min. Scale bar: 5 μm. **(D)** Percentage of cells in (C) showing cytoplasmic Nup-GFP foci. Data represent mean ±SD from at least two biological replicates. Statistical test: Two-way ANOVA with Sidak’s multiple comparison test. **p < 0.01, ns = non-significant. **(E)** Growth curves of WT and *jjj2Δ* cells. Absorbance at 600 nm is plotted over time (min). Data represent mean ±SD from four biological replicates. **(F)** Spotting assay of wild-type and *jjj2Δ* cells following a 30-minute incubation at the indicated temperatures. After heat shock, cells were spotted onto rich media (YPD) plates. **(G)** Spotting assay of wild-type and *jjj2Δ* cells harvested at stationary phase after being maintained in saturated cultures for the indicated durations. Cells were spotted onto YPD plates. **(H)** Electron micrographs of WT and *jjj2Δ* cells. Brl1-AID depleted cells (∼5.15 h auxin induction) serve as a positive control for NE herniations (yellow arrows). White arrowheads indicate mature NPCs. Note the absence of structural NE defects in *jjj2Δ* cells. Scale bar: 500 nm.

**Figure S4:**
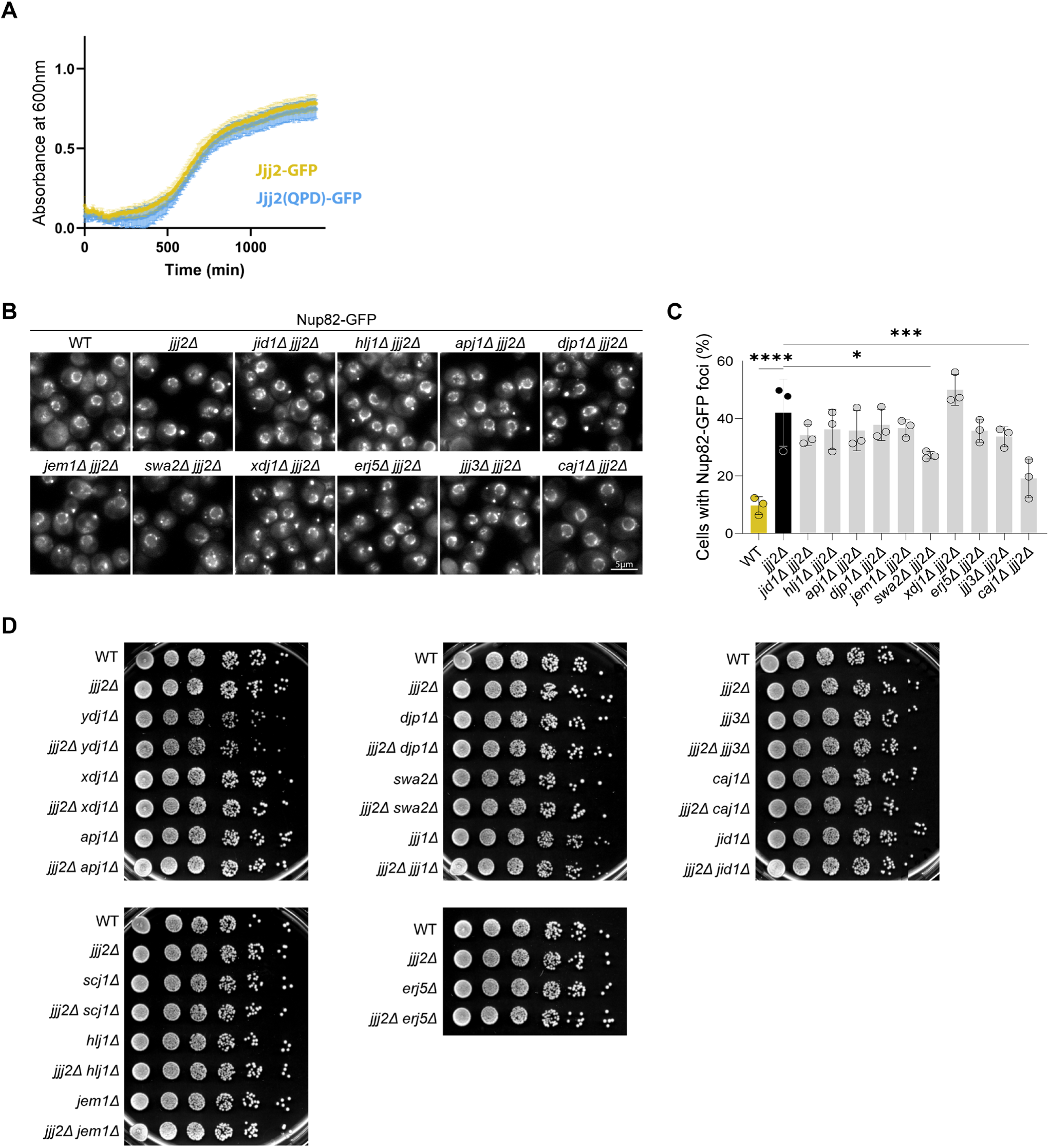
J-domain proteins do not generally regulate Nup condensation. **(A)** Growth curves of cells expressing either Jjj2-GFP or Jjj2(QPD)-GFP from the endogenous *JJJ2* genomic locus. Absorbance at 600 nm is plotted over time (min). Data represent mean ±SD from three biological replicates. **(B)** Representative fluorescence micrographs of Nup82-GFP in cells with a co-deletion of the *JJJ2* gene and the indicated non-essential J-domain protein genes. Scale bar: 5 μm. **(C)** Percentage of cells from (B) showing Nup82-GFP cytoplasmic foci. Data represent mean ±SD from three biological replicates. Statistical test: One-way ANOVA with Dunnett’s multiple comparison test. *p < 0.05, ***p < 0.001, ****p < 0.0001. **(D)** Spotting assay of cells carrying co-deletions of *JJJ2* and the indicated non-essential J-domain genes. Cells were grown on YPD medium (glucose).

**Figure S5:**
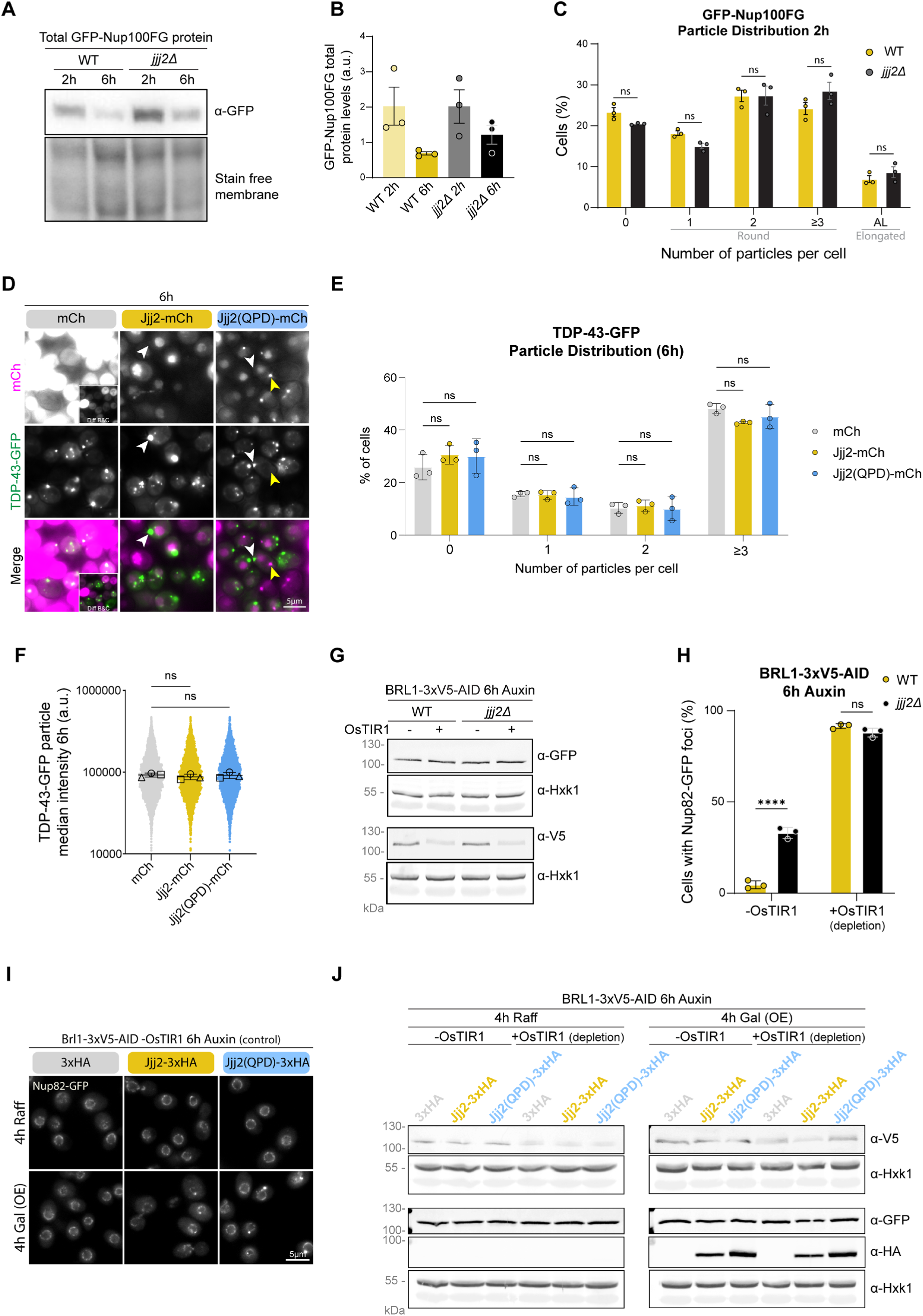
Jjj2 modulates Nup condensates. **(A)** Representative western blot showing total levels of GFP-Nup100FG in whole-cell protein extracts from WT and *jjj2Δ* cells at the indicated time points (experimental setup as in Fig. 4A). GFP-Nups were detected using an anti-GFP antibody; stain-free membrane imaging served as the loading control. **(B)** Band intensities from (A) were normalized to the total protein signal from the stain-free membrane. Data represent mean ±SD from three biological replicates. Statistical test: One-way ANOVA with Dunnett’s multiple comparison test. **(C)** Percentage of cells from the 2 h time point in Fig. 4A categorized by the number of round condensates (0, 1, 2, or ≥3) or the presence of elongated aggregate-like (AL) structures. Data represent mean ±SD from three biological replicates. Statistical test: two-way ANOVA with Sidak’s multiple comparison test. ns = non-significant. **(D)** Representative fluorescence micrographs of cells co-overexpressing either mCherry (mCh) control, Jjj2-mCh, or Jjj2(QPD)-mCh together with TDP-43-GFP (6 h time point). TDP-43-GFP forms cytoplasmic condensates (white arrowheads) that do not colocalize with Jjj2(QPD)-mCh foci (yellow arrowheads). Insets show adjusted brightness/contrast for better visualization. Scale bar: 5 μm. **(E)** Quantification of the distribution of cells in (D) categorized by the number of condensates (0, 1, 2, or ≥3). Data represent the mean ±SD from three biological replicates. Statistical test: two-way ANOVA with Dunnett’s multiple comparison test. ns = non-significant. **(F)** Superplot showing the median fluorescence intensity of TDP-43-GFP particles from (D). Individual dots represent single particles; larger black circles indicate the median of each biological replicate (n=3). Statistical test: one-way ANOVA with Dunnett’s multiple comparison test. ns = non-significant. **(G)** Western blot of whole-cell extracts from WT and *jjj2Δ* cells ±Brl1-AID depletion (from Fig. 4I). Nup82-GFP was detected with anti-GFP; Brl1-AID was detected with anti-V5. Hexokinase 1 (Hxk1) served as a loading control. **(H)** Percentage of cells from Fig. 4I showing cytoplasmic Nup82-GFP foci. Data represent mean ±SD from three biological replicates. Statistical test: Two-way ANOVA with Sidak’s multiple comparison test. ****p < 0.0001, ns = non-significant. **(I)** Representative fluorescence micrographs of Nup82-GFP in control conditions (no Brl1 depletion; -OsTIR1) from Fig. 4K, combined with overexpression (Gal) of 3xHA, Jjj2-3xHA, or Jjj2(QPD)-3xHA. Scale bar: 5 μm. **(J)** Western blot of cells from Fig. 4K and Fig. S5I. Nup82-GFP (anti-GFP), Brl1-AID (anti-V5), and 3xHA-tagged Jjj2 variants (anti-HA) were detected. Hxk1 served as a loading control.

**Figure S6:**
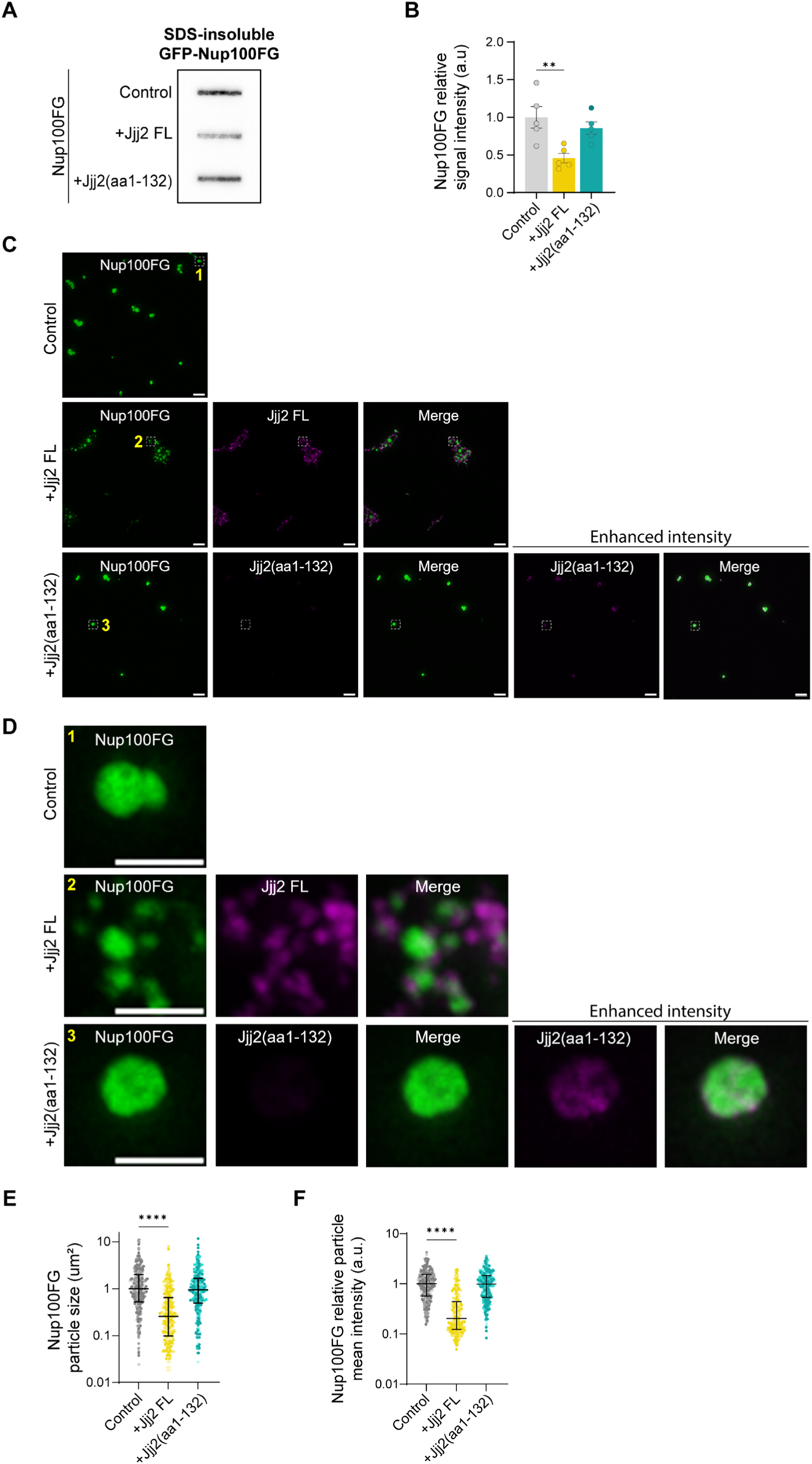
Jjj2 directly regulates Nup condensates *in vitro*. **(A)** Filter trap assay showing the 0.5% SDS-resistant aggregated fraction of purified Nup100FG. Condensates were formed in the presence of 10% PEG3350 for 1 h, either alone (control), in the presence of full-length Jjj2 (Jjj2 FL) or the Jjj2(aa1-132) truncation mutant (1:1 molar ratio). **(B)** Band intensities from (A) are expressed relative to the average intensity of the control. Data represent the mean ±SEM from five biological replicates. Statistical test: One-way ANOVA with Dunnett’s multiple comparison test. **P<0.01. **(C)** Representative fluorescence micrographs showing Nup100FG-5MF particles in the absence or presence of either Jjj2-A594 or Jjj2(aa1-132)-A594, formed in the presence of 10% PEG3350 (1 h) (molar ratio 1:1). Scale bar: 1 μm. A594 (Alexa Fluor™ 594 C5 Maleimide), 5FM (fluorescein-5-maleimide). **(D)** Representative micrographs of the insets indicated by yellow numbers in (C). Scale bar: 1 μm **(E-F)** Quantification of particle size (E) and mean fluorescence intensity (F) of Nup100FG-5MF particles from (D). Intensity values are normalized to the median of the control. Graphs show the median ± interquartile range of 300 particles per condition (n=3 independent replicates, 100 particles each). A color gradient highlights data points belonging to each independent replicate. Statistics: Kruskal-Wallis test with Dunn’s multiple comparison. ∗∗∗∗p < 0.0001.

**Figure S7:**
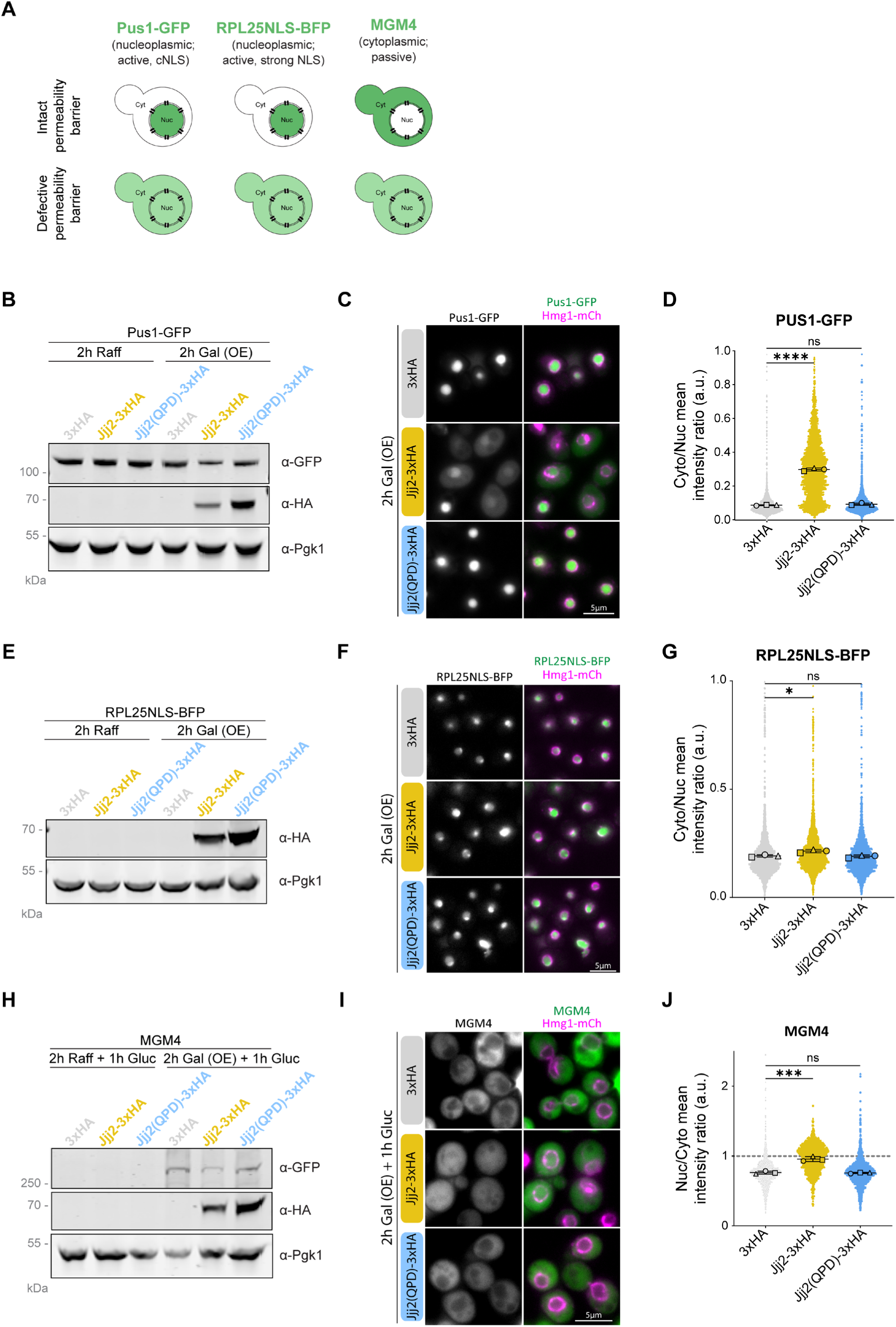
Jjj2 overexpression leads to permeability barrier defects. **(A)** Schematic of the reporters used to assess the permeability barrier defects (Pus1-GFP, RPL25NLS-BFP and MGM4) and their expected subcellular localization in conditions with intact or defective permeability barrier. **(B)** Western blot of Pus1-GFP-expressing cells upon overexpression (Gal) of 3xHA, Jjj2-3xHA, or Jjj2(QPD)-3xHA. Raff is used as a non-overexpression control. Pus1-GFP (anti-GFP) and 3xHA-tagged Jjj2 variants (anti-HA) were detected. Pgk1 served as a loading control. **(C)** Representative fluorescence micrographs of Pus1-GFP expressing cells upon 2 h of overexpression (Gal) of 3xHA, Jjj2-3xHA, or Jjj2(QPD)-3xHA. Scale bar: 5 μm. **(D)** Superplot showing the Pus1-GFP cytoplasmic/nucleoplasmic mean intensity ratio from cells in (C). Small colored shapes represent individual cells; larger black shapes indicate the mean of each biological replicate (n=3). Statistical test: one-way ANOVA with Dunnett’s multiple comparison test. ****p < 0.0001. **(E)** Western blot of RPL25NLS-BFP expressing cells upon overexpression (Gal) of 3xHA, Jjj2-3xHA, or Jjj2(QPD)-3xHA. Raff is used as a non-overexpression control. 3xHA-tagged Jjj2 variants (anti-HA) were detected. Pgk1 served as a loading control. **(F)** Representative fluorescence micrographs of RPL25NLS-BFP expressing cells upon 2h of overexpression (Gal) of 3xHA, Jjj2-3xHA, or Jjj2(QPD)-3xHA. Scale bar: 5 μm. **(G)** Superplot showing the RPL25NLS-BFP cytoplasmic/nucleoplasmic mean intensity ratio from cells in (F). Small dots represent individual cells; larger black shapes indicate the mean of each biological replicate (n=3). Statistical test: one-way ANOVA with Dunnett’s multiple comparison test. *p < 0.05. **(H)** Western blot of cells upon inducible overexpression (Gal) of MGM4 together with 3xHA, Jjj2-3xHA, or Jjj2(QPD)-3xHA. Raff is used as a non-overexpression control. Glucose was added for 1 h to stop overexpression and to allow newly synthesized proteins to mature and equilibrate as in (Popken et al., 2015). MGM4 (anti-GFP) and 3xHA-tagged Jjj2 variants (anti-HA) were detected. Pgk1 served as a loading control. **(I)** Representative fluorescence micrographs of cells upon inducible overexpression (Gal) of MGM4 together with 3xHA, Jjj2-3xHA, or Jjj2(QPD)-3xHA. Glucose was added for 1 h to stop overexpression and to allow newly synthesized MGM4 proteins to mature and equilibrate as in (Popken et al., 2015). Scale bar: 5 μm. **(J)** Superplot showing the MGM4 nucleoplasmic/cytoplasmic mean intensity ratio from cells in (I). Small colored shapes represent individual cells; larger black shapes indicate the mean of each biological replicate (n=3). Statistical test: one-way ANOVA with Dunnett’s multiple comparison test. **p < 0.01.

**Figure S8:**
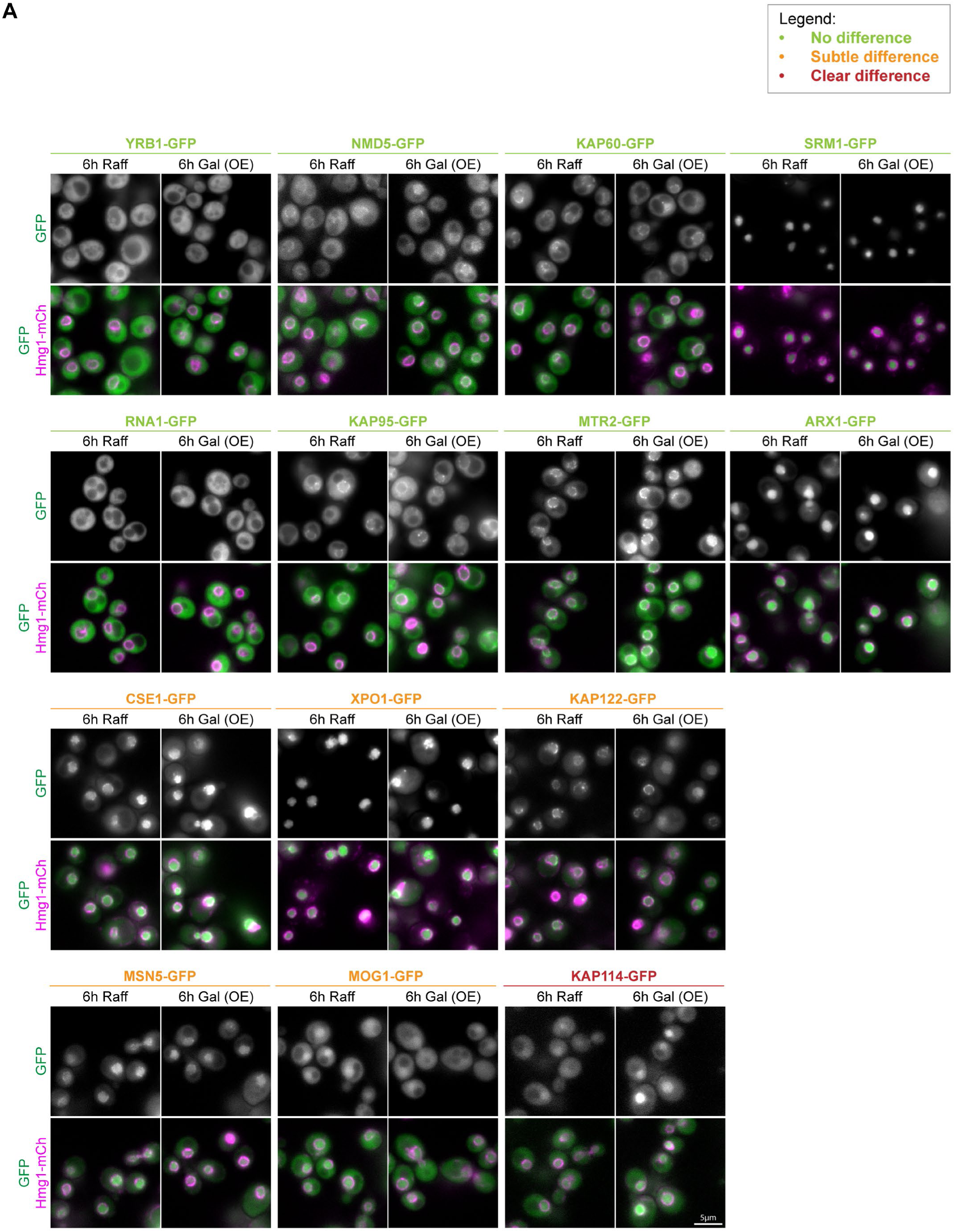
Jjj2 overexpression does not generally impact nuclear transport receptor subcellular localization. **(A)** Representative fluorescence micrographs of cells expressing the indicated GFP-tagged nuclear transport receptors (NTRs) following 6 h of overexpression (Gal) of either 3xHA (control) or Jjj2-3xHA. Raffinose (Raff) serves as a non-overexpression control. NTRs are categorized based on their localization pattern: the majority of NTRs maintain normal subcellular distribution (labeled in green); a subset exhibits subtle changes (labeled in orange); and a few show clear differences (labeled in red). Scale bar: 5 μm.

**Figure S9:**
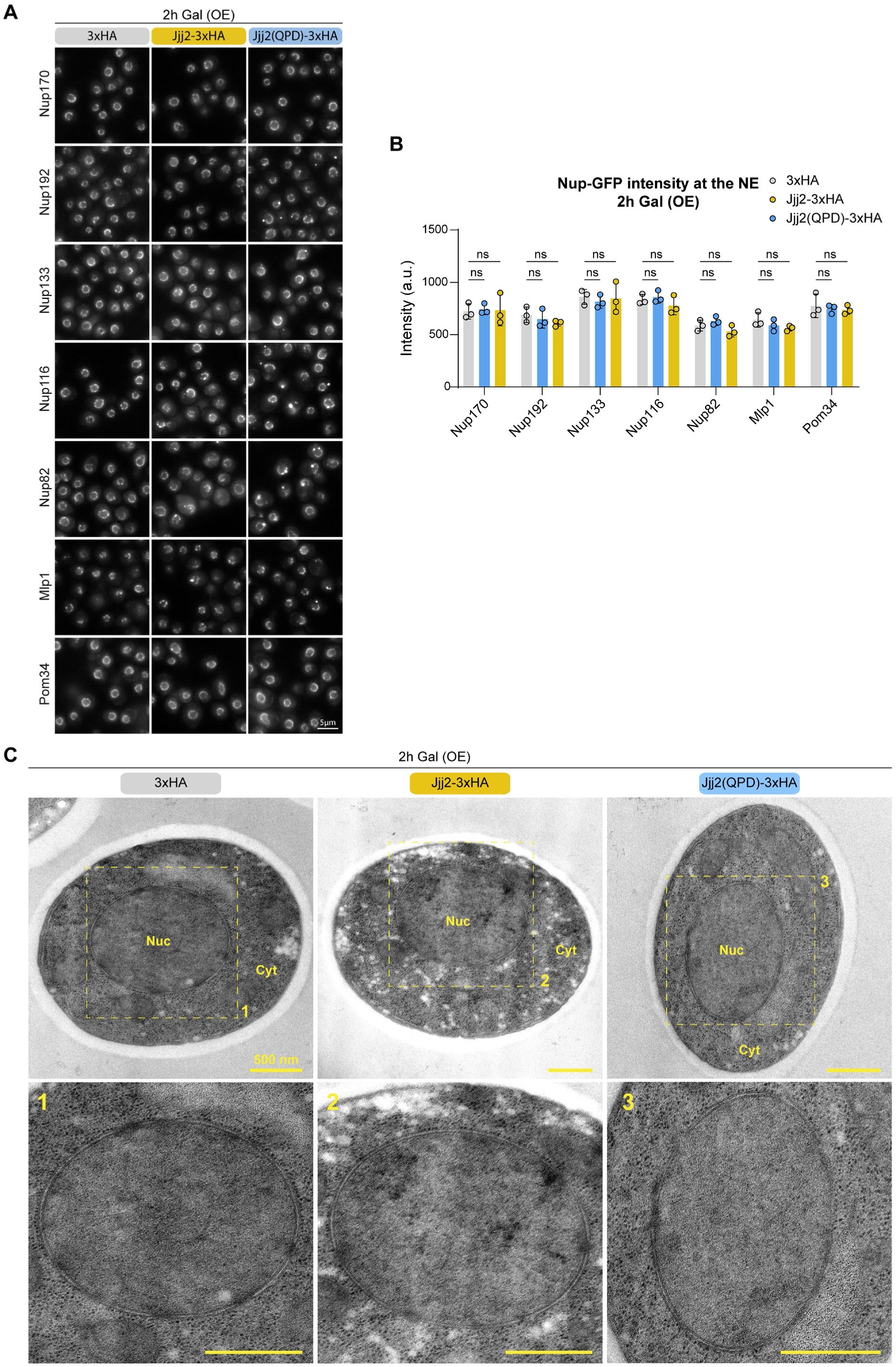
Jjj2 overexpression does not affect Nup levels at the nuclear rim or nuclear envelope structure. **(A)** Representative fluorescence micrographs of cells expressing the indicated GFP-tagged Nups following 2 h of overexpression (Gal) of either 3xHA (control), Jjj2-3xHA or Jjj2(QPD)-3xHA. Scale bar: 5 μm. **(B)** Quantification of Nup-GFP fluorescence intensity at the nuclear envelope (NE) from cells in (A). Data represent mean ±SD from at least three biological replicates. Statistical test: Two-way ANOVA with Dunnett’s multiple comparison test. ns = non-significant. **(C)** Electron micrographs of cells following 2 h of overexpression (Gal) of either 3xHA (control), Jjj2-3xHA or Jjj2(QPD)-3xHA. Note the absence of obvious structural NE defects. Scale bar: 500 nm.

